# *VPS41* recessive mutation causes ataxia and dystonia with retinal dystrophy and mental retardation by inhibiting HOPS function and mTORC1 signaling

**DOI:** 10.1101/2019.12.18.867333

**Authors:** R.E.N. van der Welle, R. Jobling, C. Burns, P. Sanza, C. ten Brink, A. Fasano, L. Chen, F.J. Zwartkruis, S. Zwakenberg, E.F. Griffin, J. van der Beek, T. Veenendaal, N. Liv, S. Blaser, C. Sepulveda, A.M. Lozano, G. Yoon, C.S. Asensio, G.A. Caldwell, K.A. Caldwell, D. Chitayat, J. Klumperman

## Abstract

The vacuolar protein sorting protein 41 (VPS41) is a neuroprotective protein in models of Parkinson’s disease (PD). As part of the HOPS (Homotypic fusion and Protein Sorting) complex, VPS41 regulates fusion of lysosomes with late endosomes and autophagosomes. Independent of HOPS, VPS41 regulates transport of newly synthesized lysosomal membrane proteins and secretory proteins. Here we report two brothers with compound heterozygous mutations in *VPS41* (*VPS41*^*R662\**^ and *VPS41*^*S285P*^), born to healthy and non-consanguineous parents. Both patients displayed transient retinal dystrophy, ataxia and dystonia, with brain MRI findings of cerebellar atrophy and a thin saber-shape corpus callosum. Patient-derived fibroblasts contained enzymatically active lysosomes that were poorly reached by endocytic cargo and failed to attract the mTORC1 complex. Consequently, transcription factor TFE3, a driver of autophagy and lysosomal genes, showed continuous nuclear localization which resulted in elevated LC3-II levels and an impaired response to nutrient starvation. CRISPR/CAS *VPS41* HeLa knockout cells showed a similar phenotype that could be rescued by wildtype VPS41 but not by VPS41^S285P^ or VPS41^R662*^. mTORC1 inhibition was also seen after knockout of HOPS subunits VPS11 or VPS18. Regulated neuropeptide secretion in PC12 *VPS41* knockout cells was rescued by VPS41^S285P^ expression, indicating that this HOPS-independent function was preserved. Co-expression of the VPS41^S285P^ and VPS41^R662*^ variants in a *C. elegans* model of PD abolished the protective effect of VPS41 against α-synuclein-induced neurodegeneration. We conclude that both disease-associated *VPS41* variants specifically abrogate HOPS function, which leads to a delay in endocytic cargo delivery to lysosomes, mTORC1 inhibition and irresponsiveness to autophagic clues. Our studies signify a link between HOPS function and mTORC1 signaling and imply that HOPS function is required for the neuroprotective effect of VPS41 in PD.

## Introduction

While lysosomes are responsible for the degradation and recycling of intra- and extracellular substrates(1), they are increasingly recognized as regulators of cellular homeostasis and key players in nutrient sensing and transcriptional regulation(2–5). Hence, the digestive properties of lysosomes give them a major role in the control of cellular metabolism and nutrient homeostasis. To accomplish this, HOPS (Homotypic Fusion and Protein Sorting) regulates fusion of lysosomes with endosomes and autophagosomes(6–11).

Vacuolar Protein Sorting 41 (VPS41) (Chr7p14.1), a defining component of HOPS, is a 100 kD protein that contains a WD40, TRP-like (tetratricopeptide repeat), CHCR (Clathrin Heavy Chain Repeat) and RING (Really Interesting New Gene)-H2 Zinc Finger domain(12,13). *VPS41* knockout mice die early in utero, defining VPS41 as an essential protein for embryonic development(14). HOPS-associated VPS41 is recruited to endosomes by binding to Rab7 and its interactor protein RILP and to lysosomes by interacting with Arf-like protein 8b (Arl8b)(15,16). Depletion of VPS41 impairs HOPS dependent delivery of endocytic cargo to lysosomes and causes a defect in autophagosomal flux(6,17). Independent of HOPS, VPS41 is required for transport of lysosomal membrane proteins from the *trans*-Golgi-Network (TGN) to lysosomes(18),(19), a pathway known as the Alkaline Phosphatase (ALP) pathway in yeast(20–24). Furthermore, in secretory cells and neurons, VPS41 is required for regulated secretion of neuropeptides(25–31). Both in yeast and in secretory cells it was suggested that VPS41 forms a coat on adaptor protein 3 (AP-3) containing membranes that exit the TGN(23,24,31). Together these data show that VPS41, as part of HOPS, is required for lysosomal fusion events and, independent of HOPS, for transport of lysosomal membrane proteins and regulated secretion.

Interestingly, *VPS41* overexpression protects dopaminergic neurons against neurodegeneration in a transgenic *C. elegans* model of Parkinson’s Disease (PD), in which α-synuclein is locally overexpressed (32–34). Similarly, when *VPS41* is overexpressed in human neuroglioma cells, α-synuclein protein levels are reduced(32). This neuroprotective effect requires interactions of VPS41 with Rab7 and AP-3(35). Recent studies using a *C. elegans* model for Alzheimer’s disease showed that overexpression of human *VPS41* also mitigates Aβ-induced neurodegeneration of glutamatergic neurons. This requires Arl8 activity rather than Rab7 and AP3, indicating that VPS41 through different interaction partners can trigger divergent neuroprotective mechanisms(35). However, how VPS41 function leads to neuroprotection remains to be elucidated.

Naturally occurring SNPs in *VPS41* have been described (T52R, T146P and A187T)(32,36), but thus far no patients bearing mutations in *VPS41* were identified. In this study, we present brothers with severe neurological features (e.g. transient retinal dystrophy, ataxia and dystonia accompanied by retinal dystrophy and mental retardation with brain MRI findings of cerebellar atrophy and thin corpus callosum) of unknown aetiology. Exome sequencing showed that both patients were compound heterozygous for variants in VPS41 [NM_014396.19: c.853T>C, NP_055211.2: pSer285Pro (S285P); NM_014396.6: c.1984C>T, NP_055211.2: pArg662Stop (R662*)]. To our best knowledge, this is the first report of patients bearing biallelic *VPS41* variants displaying neurological manifestations. We show that the *VPS41* variants in patient cells prevents formation of the HOPS complex, leading to a defect in the delivery of endocytosed and autophagic cargo to lysosomes, inhibition of the mTORC1 complex, and a constitutive upregulation of autophagy. This renders cells unable to respond to changes in nutrient conditions and abolishes the neuroprotective effect of VPS41 in a *C. elegans* model of PD.

## Results

### Clinical presentation of two patients with biallelic mutations in VPS41

Two brothers, born to healthy and non-consanguineous parents, were noted to be hypotonic shortly after birth. In infancy they showed global developmental delay, low muscle tone, and marked tremor which further impaired their fine motor skills. Both brothers showed absent deep tendon reflexes (DTRs) and their plantars were extensor. Both patients developed upper extremity tremor and significant lower limbs’ ataxia and spastic ataxia. Brain MRI in both cases showed mild hypomyelination (Figure 1A, B). Brain MRI studies of the younger brother (patient 2) revealed mild hypoplasia of the cerebellum and cerebellar vermis and a thin corpus callosum. A repeat brain MRI at 10 years of age confirmed the thin corpus callosum, mild progression of the cerebellar atrophy, bilateral hypointensity in the globus pallidus (GP) compatible with early iron deposition, as well as abnormal hyperintensity in the dentate nucleus and cerebellar cortex (Figure 1B). Analyses of genes known to cause X-linked mental retardation and dystonia as well as genes associated with iron deposition in the basal ganglia were tested but no mutations were found (Table S1). Whole exome sequencing performed on both patients revealed compound heterozygote variants in *VPS41*.

**Figure 1.**
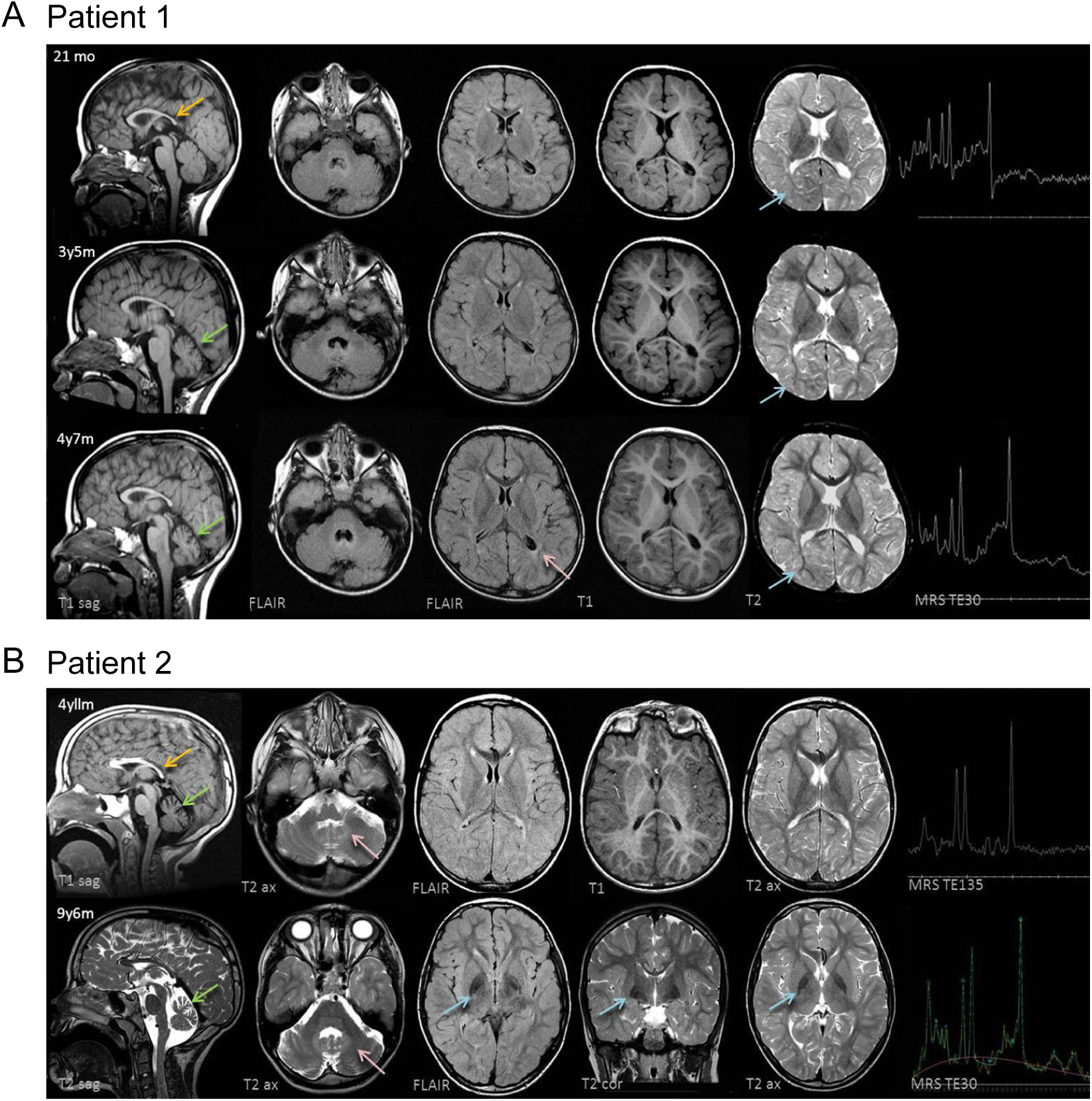
Mutations in *VPS41* result in the development of a neurodegenerative disease. **(A)** Patient 1 (older sibling) underwent 3 MRI studies. Top row: 21 months; Middle row: 3 years, 5 months; Bottom row: 4 years 7 months. The corpus callosum is thin with a saber-shape (yellow arrow) configuration (splenium more thinned than the genu and body). The shape remains consistent over time. The vermis is normal in configuration and size initially, but demonstrates volume loss on follow-up examinations (green arrows). There is delay in myelin maturation present on T1 and T2 weighted axial images (blue arrow on T2 shows lack of peripheral myelin (arborization) on initial examination. There is further myelin development over time (blue arrows), although myelin maturation is slow. The TE30 MRS is age appropriate at both 21 months of age and 4 years 7 months of age. By 4 years 7 months of age, peritrigonal gliosis (FLAIR, pink arrow) and volume loss is obvious. **(B)** Patient 2 (younger sibling) underwent 2 MRI studies. Top row: 4 years, 11 months; Bottom row: 9 years, 6 months. The corpus callosum is thin with a saber-shape (yellow arrow) configuration (splenium more thinned than the genu and body). The shape remains consistent over time. The vermis demonstrates volume loss on both examinations (green arrows). The dentate nuclei (pink arrows) are bright on both examinations. There is cerebellar hemispheric volume loss on both studies. Myelin maturation is minimally delayed on T2 weighted image at 4 years 11 months of age, but normal on follow up examination. The TE 144 MR Spectroscopy (MRS) and follow-up TE 30 MRS are both age appropriate. There is no peritrigonal gliosis, although there is slightly less white matter in the peritrigonal region than expected for age. There is iron deposition in the basal ganglia on FLAIR and T2 weighted images at 9 years 8 months of age (blue arrows). Iron deposition was also present in the subthalamic nuclei (not shown).

Each of the parents was found to be heterozygote for a variant in the *VPS41* gene; the father carries a point mutation in the WD40 domain (NM_014396.19: c.853T>C, NP_055211.2: pSer285Pro, hereafter referred to as VPS41^S285P^), the mother carries a nonsense mutation in the Clathrin Heavy Chain Repeat (CHCR) resulting in a truncated protein (NM_014396.6: c.1984C>T, NP_055211.2: pArg662Stop; hereafter referred to as VPS41^R662*^) (Figure S1).

### Patient fibroblasts and *VPS41*^*KO*^ cells contain small-sized, acidified lysosomes active for cathepsin B

To study the cellular effects of the VPS41 variants we obtained skin biopsies from the youngest brother (case 2) (*VPS41*^*S285P/R662\**^), an independent control (*VPS41*^*WT/WT*^), the patients’ father (*VPS41*^*WT/S285P*^) and the patients’ mother (*VPS41*^*WT/R662\**^) and made primary fibroblast cultures. Since the older brother was found to have a de novo duplication at Chr1q21.1 of unknown significance (Supplemental information), only fibroblasts of the second patients were obtained. First we studied the effects of the mutations on lysosomal biogenesis and function. Electron microscopy (EM) showed a high variation in appearance of endo-lysosomal compartments in primary fibroblasts (Figure S2), but no swollen lysosomes in the patient-derived cells. This indicates that the patient-specific mutations in *VPS41* do not cause a classical lysosomal storage phenotype(37). To study lysosomal acidification and hydrolase activity we performed fluorescence microscopy using lysotracker-Green and MagicRed-cathepsin B. This showed an increase in the number of acidic and cathepsin B-active puncta in patient fibroblasts as compared to control or parental cells (Figure S3A, quantified in S3A’). Concomitantly, immunofluorescence of the lysosomal membrane protein LAMP-1 showed twice as much puncta in patient cells (Figure 2A, quantified in 2A’). A similar phenotype was found in HeLa^*VPS41KO*^ cells, i.e. Hela cells knockout for *VPS41* using CRISPR/Cas9 methodology (Figures 2B and C, quantified in 2C’ and Figure S3B, quantified in S3B’). LAMP-1 resides in lysosomes as well as late endosomes, which by EM are distinguished by the presence or absence of degraded material, respectively(38–41). By immuno-EM we found that *VPS41*^*S285P/R662\**^ fibroblasts contain numerous LAMP-1 positive late endosomes and lysosomes which are positive for cathepsin B (Figures 2D and Figure S3C), but that the lysosomes are significantly smaller than in *VPS41*^*WT/WT*^, *VPS41*^*WT/S285P*^ or *VPS41*^*WT/R662\**^ fibroblasts (Figure 2E). The relative labeling density of LAMP-1 in the lysosomal membrane remained equivalent to control cells (Figure 2F), but since lysosomes were smaller the absolute number of gold particles per lysosome had decreased (Figure 2G). Similar data were obtained for LAMP-2 (Figure S4A and B). We conclude from these data that the biallelic VPS41^R662*^ and VPS41^S285P^ variants induce the presence of small-sized, enzymatically-active lysosomes, which at steady state conditions contain normal LAMP levels.

**Figure 2.**
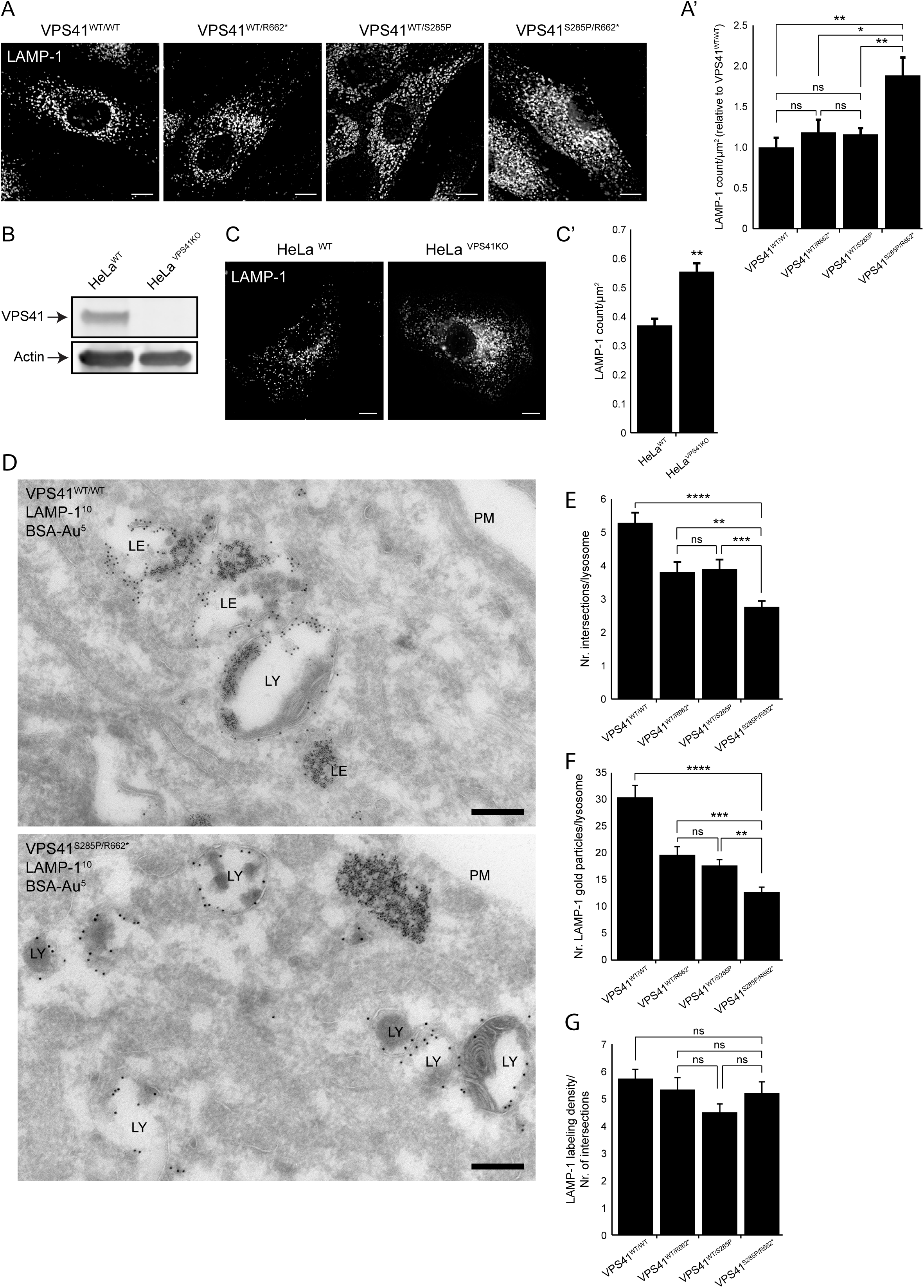
VPS41 patient and HeLa *VPS41* knockout cells contain more but smaller lysosomes. **(A)** Immuno-fluorescence microscopy of control, parental and patient fibroblasts labeled for LAMP-1. Patient (*VPS41*^*S285P/R662\**^) fibroblasts show more LAMP-1 puncta, which are distributed throughout the cell. **(A’)** Quantification shows a significant increase in the number of LAMP-1 positive compartments. Based on >15 cells per condition (n=3). Bars 10µm. Error bars represent Standard Error of the Mean (SEM). *p<0.05, ***p<10^−4^; one-ANOVA with Tukey’s correction for multiple comparisons. **(B)** HeLa VPS41 knockout (HeLa^*VPS41KO*^) cells made using CRISPR/Cas9 methodology. Western Blot analysis confirms a full knockout. **(C)** LAMP-1 immunofluorescence of HeLa^*VPS41KO*^ cells. Similar to patient-derived fibroblasts more LAMP-1 positive compartments are seen (quantified in **C’**). Based on >10 cells per cell line per experiment (n=3). Bars 10µm. Error bars represent the SEM. Unpaired *t* test, **p<0.01. (Figure S3). **(D)** Immuno-electron microscopy of *VPS41*^*WT/WT*^, *VPS41*^*WT/S285P*^, *VPS41*^*WT/R662\**^ and *VPS41*^*S285P/R662\**^ fibroblasts loaded for 2h with BSA conjugated to 5 nm gold (BSA-Au^5^) and labeled for LAMP-1 (10 nm gold particles). Lysosomes are recognized by presence of degraded, electron-dense material. PM= plasma membrane, LE= late endosome, LY= lysosome. Bar is 200nm. **(E)** Morphometrical analysis shows that lysosomes in patient fibroblasts are significantly smaller than in control and parental cells. Quantification based on >100 randomly selected lysosomes per condition. Error bars represent the SEM. **p<0.01, ****p<10^−5^; one-ANOVA with Tukey’s correction for multiple comparisons. **(F)** Quantitation of LAMP-1 gold particles per lysosome. *VPS41*^*S285P/R662\**^ lysosomes have significantly less LAMP-1 compared to control and parental cells (n>46 lysosomes per condition). Similar results were obtained for LAMP-2 (Figure S4). Error bars represent SEM. **p<0.01, ***p<10^−4^, ****p<10^−5^; one-ANOVA with Tukey’s correction for multiple comparisons. **(G)** Relative labeling density of LAMP-1. Number of LAMP-1 gold particles per lysosome was divided by the number of grid intersections representing lysosomal size. No significant difference was found between *VPS41*^*WT/WT*^, *VPS41*^*WT/S285P*^, *VPS41*^*WT/R662\**^ or *VPS41*^*S285P/R662\**^. Error bars represent the SEM; one-ANOVA with Tukey’s correction for multiple comparisons.

### VPS41^R662*^ poorly interacts with other HOPS subunits

To understand the effects of the VPS41 mutations at the molecular level, we investigated the ability of VPS41^R662*^ and VPS41^S285P^ to interact with other subunits of the HOPS complex. Hereto we performed co-immunoprecipitation (co-IP) studies in HeLa cells expressing GFP-tagged constructs of VPS41^WT^, VPS41^S285P^ or VPS41^R662*^ and respectively FLAG- and HA-tagged constructs of VPS18 or VPS33A. Expression of all constructs was readily detectable by Western blot (Figure 3A and S5). The premature stop codon in VPS41^R662*^ resulted in a truncated protein of lower molecular weight. VPS41^S285P^ interacted with both VPS18 and VPS33A, but interaction of VPS41^R662*^ with the other HOPS subunits was strongly reduced (Figure 3A and S5). Since interaction between VPS41 and VPS18 requires the presence of the C-terminal RING domains that is absent from VPS41^R662*^ (Figure S1), these data are in agreement with present literature(42).

**Figure 3.**
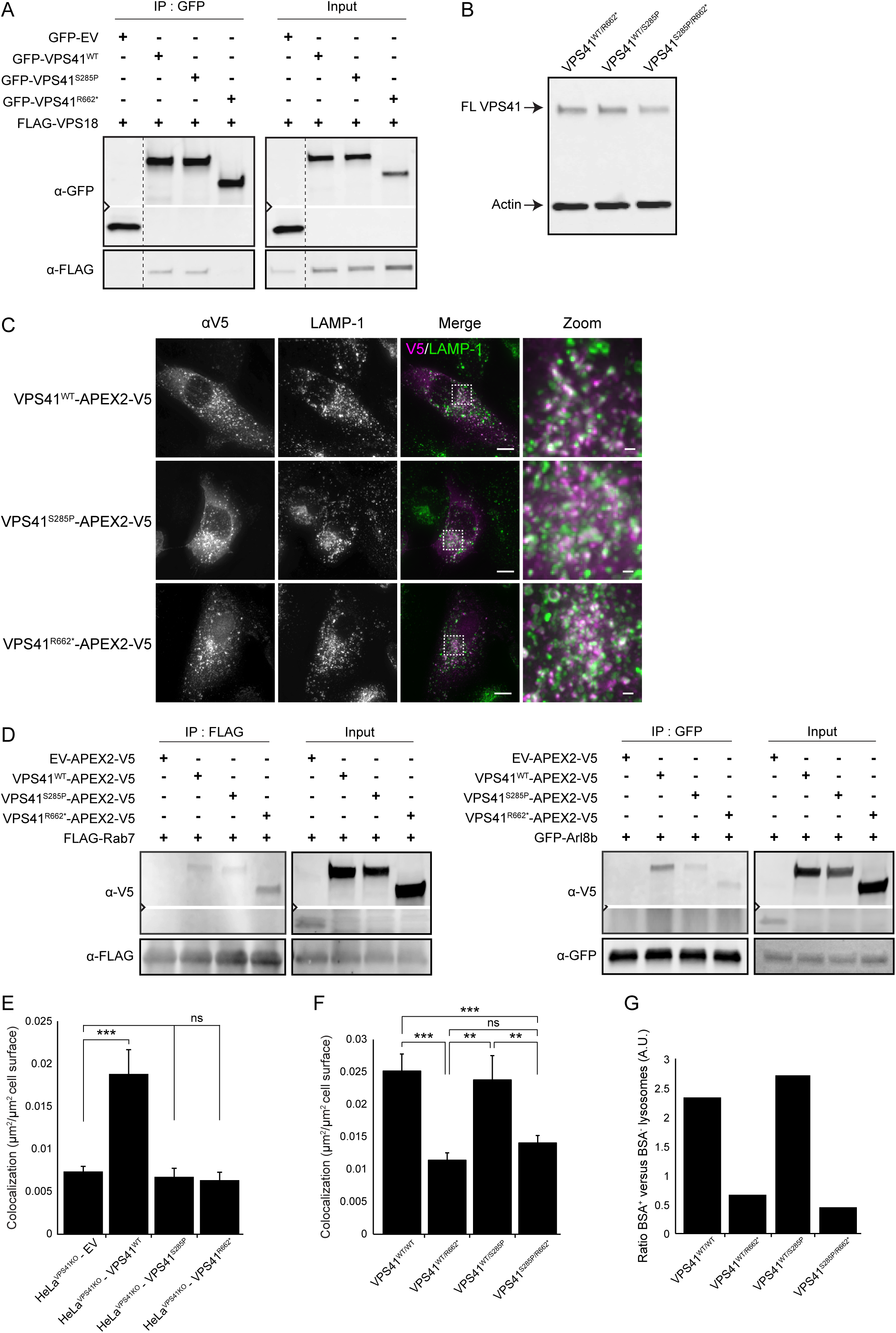
Loss of VPS41^WT^ results in impaired HOPS dependent late endosome – lysosome fusion. **(A)** Immunoprecipitation (IP) on HeLa cells co-expressing GFP-empty vector (EV), GFP-VPS41^WT^, GFP-VPS41^S285P^ or GFP-VPS41^R662*^ and FLAG-VPS18. GFP-VPS41^WT^ and GFP-VPS41^S285P^ show interaction with FLAG-VPS18. Interaction between GFP-VPS41^R662*^ and FLAG-VPS18 is almost completely lost. Note that the stopcodon in *VPS41*^*R662\**^ leads to a truncated protein of lower MW. n=3. Similar results were obtained for HA-VPS33A (Figure S5). **(B)** Western Blot of primary fibroblasts derived from mother (*VPS41*^*WT/R662\**^), father (*VPS41*^*WT/S285P*^) or patient (*VPS41*^*S285P/R662\**^). Full-length VPS41 (FL VPS41) representing VPS41^WT^ and VPS41^S285P^ is observed in all cells. Truncated VPS41^R662**\***^ expressed in patient and maternal fibroblasts is not detected. **(C)** HeLa^*VPS41KO*^ cells transfected with VPS41^WT^-APEX2-V5, VPS41^S285P^-APEX2-V5 or VPS41^R662*^-APEX2-V5 constructs were labeled for LAMP-1 and V5 for immunofluorescence microscopy. All VPS41 variants co-localized with LAMP-1, indicating that VPS41^S285P^ and VPS41^R662*^ are recruited to late endosomes/lysosomes. Bars 10µm, zoom of squared area 1µm. **(D)** IP on Hela cells co-expressing VPS41^WT^-APEX2-V5, VPS41^S285P^-APEX2-V5 or VPS41^R662*^-APEX2-V5 and FLAG-RAB7 or GFP-Arl8b. The IP was performed on FLAG or GFP respectively. All VPS41 variants interact with Rab7 and Arl8b (n=3). **(E)** Rescue experiments. HeLa^*VPS41KO*^ cells transfected with VPS41^WT^-APEX2-V5 showed a significant increase in colocalization between Dextran and SiR-Lysosome cathepsin D, indicating rescue of the endocytosis phenotype. Neither *VPS41* variant could rescue HOPS complex functionality (Figure S6A) (n=3). Error bars represent the SEM. ***p<10^−4^; one-ANOVA with Bonferroni correction for multiple comparisons. **(F)** *VPS41*^*WT/WT*^, VPS41^WT/S285P^. *VPS41*^*WT/R662\**^ and *VPS41*^*S285P/R662\**^ primary fibroblasts were incubated with Dextran-ALEXA568 and SiR-Lysosome cathepsin D for 2h and 3h respectively. Co-localization representing delivery of Dextran to enzymatically active lysosomes was decreased in *VPS41*^*WT/R662\**^ and *VPS41*^*S285P/R662\**^ cells, indicating reduced fusion efficiency between late endosomes and lysosomes (Figure S6B) (n=3). Error bars represent the SEM. **p<0.01, ***p<10^−4^; one-ANOVA with Tukey’s correction for multiple comparisons. **(G)** *VPS41*^*WT/WT*^, *VPS41*^*WT/S285P*^, *VPS41*^*WT/R662\**^ and *VPS41*^*S285P/R662\**^ fibroblasts were incubated with BSA-Au^5^ for 2h and labeled for LAMP-1 (10 nm gold particles) for immuno-EM (Figure 2D). LAMP-1-positive lysosomes were scored positive or negative for BSA-Au^5^ and the ratio between BSA^+^ and BSA^-^ lysosomes was calculated. Both *VPS41*^*WT/R662\**^ and *VPS41*^*S285P/R662\**^ show a strong decrease in BSA^+^-lysosomes indicating a fusion defect between late endosomes and lysosomes.

Truncated proteins are often prematurely degraded. To test this for VPS41^R662*^, we analyzed the expression levels of the *VPS41* variants in primary fibroblasts by Western Blot. Full length VPS41, representing both VPS41^WT^ and VPS41^S285P^, was readily detectable. However, no VPS41^R662*^ was seen in either VPS41^WT/R662*^ maternal or VPS41^S285P/R662*^ patient fibroblasts (Figure 3B). Since the antibody does recognize truncated VPS41^R662*^ when overexpressed (Figure 3A and S5), we conclude that VPS41^R662*^ protein levels are below detection in primary fibroblasts derived from the patient and mother. Of note, VPS41 expression levels are higher in a.o. neuronal cell types (https://www.proteinatlas.org/ENSG00000006715-VPS41/), which renders it possible that other cell types in the patients express residual levels of VPS41^R662*^.

Together these experiments show that VPS41^R662*^ is unable to bind other HOPS components whereas VPS41^S285P^ binds VPS18 and VPS33A. Also, VPS41^R662*^ shows an overall low expression level.

### VPS41^S285P^ and VPS41^R662*^ both cause a defect in late endosome – lysosome fusion

Depletion of VPS41 by RNAi results in a decrease in HOPS-dependent fusion between late endosomes and lysosomes(6,19). To establish the effect of the individual VPS41 mutants on HOPS function, we transiently transfected HeLa^*VPS41KO*^ cells with APEX2-V5 tagged constructs of VPS41^WT^, VPS41^S285P^ or VPS41^R662*^. The subcellular distribution of these constructs was explored by immunofluorescence microscopy. VPS41^WT^ was found dispersed in the cytoplasm as well as in distinct fluorescent puncta that co-localized with LAMP-1, consistent with previous studies(16,43) (Figure 3C). Remarkably, both VPS41^S285P^ and VPS41^R662*^ showed a similar distribution as VPS41^WT^, despite the lack of the C-terminus in VPS41^R662*^. To address the mechanism of membrane association of VPS41 we performed co-IP’s using APEX-V5-tagged VPS41 constructs and FLAG and GFP-tagged constructs of Arl8b and Rab7. This showed that VPS41^R662*^ binds Rab7 and Arl8 to the same extent as VPS41^WT^ and VPS41^S285P^ (Figure 3D). These data indicate that Rab7 and Arl8b can recruit VPS41^R662*^ to late endosomes and lysosomes independent of its incorporation in the HOPS complex or presence of the C-terminus.

To test whether either of the mutants allows for HOPS function, *VPS41*^*KO*^ cells were incubated for 2 hours with the endocytic marker Dextran-ALEXA568 and SiR-Lysosome to mark active cathepsin D compartments. Co-localization between these probes indicates the transfer of endocytosed cargo to enzymatically active lysosomes. *VPS41*^*KO*^ cells transfected with Empty Vector (EV) showed low levels of co-localization (Figure 3E and Figure S6A). Transfection with VPS41^WT^ increased co-localization, indicating a restoration of HOPS function (Figure S6A). Transfection with VPS41^R662*^ did not rescue the endocytosis defect, which was expected since VPS41^R662*^ fails to bind other HOPS components (Figure 3A). Surprisingly, however, VPS41^S285P^ also failed to rescue the endocytosis defect (Figure 3E and Figure S6). Thus, even though VPS41^S285P^ binds VPS18 and VPS33A (Figure 3A and Figure S5) and localizes to endo-lysosomes (Figure 3C) it cannot form a functional HOPS complex. A plausible explanation is misfolding of the protein due to the Arginine to Proline substitution, which is predicted to cause a conformation change (http://genetics.bwh.harvard.edu/pph2/)(44). Concluding, these data indicate that neither VPS41 variant allows formation of a functional HOPS complex.

### Patient fibroblasts are compromised in delivery of endocytosed cargo to enzymatically active lysosomes

To assess the process of late endosome – lysosome fusion in patient cells we applied the Dextran-ALEXA568 and SiR-Lysosome cathepsin D colocalization assay to the primary fibroblast cultures. This confirmed an endocytosis defect in *VPS41*^*S285P/R662\**^ (patient) fibroblasts. Notably, however, also the maternal *VPS41*^*WT/R662\**^ cells showed a decreased co-localization of Dextran-ALEXA568 and SiR-Lysosome cathepsin D. By contrast, *VPS41*^*WT/S285P*^ (paternal) fibroblasts were not affected in endocytosis (Figure 3F and Figure S6B). To investigate the block in endocytosis at the EM level, we incubated primary fibroblasts for 2 hours with BSA conjugated to 5 nm gold particles (BSA-Au^5^) and processed cells for immuno-EM. Sections were labeled for LAMP-1 or LAMP-2 and randomly screened for LAMP-positive lysosomes(40) (Figure 2D), which were scored positive or negative for BSA-Au^5^. This showed a strong decrease in the delivery of BSA-Au^5^ to LAMP-positive lysosomes of patient fibroblasts (Figure 3G). In agreement with the fluorescent data the maternal *VPS41*^*WT/R662\**^ fibroblasts also showed a decrease in BSA-Au^5^-positive lysosomes, whereas the paternal *VPS41*^*WT/S285P*^ fibroblasts were not affected. Together these data show that in patient fibroblasts the delivery of endocytosed cargo to enzymatically active lysosomes is delayed, which is explained by that neither VPS41 variant allows formation of a functional HOPS complex (Figure 3E). The endocytic delay observed in the maternal fibroblasts, indicates that in this case the defect is not fully compensated for by the VPS41^WT^ allele.

### Loss of VPS41^WT^ causes a defect in autophagic response to starvation

In addition to late endosome-lysosome fusion, the HOPS complex is required for fusion of autophagosomes with lysosomes(45–47). A block in autophagosome-lysosome fusion is predicted to result in increased numbers of autophagosomes containing lipidated LC3 (LC3-II). We addressed LC3-II levels in the primary fibroblast cultures by Western blotting. In nutrient-rich conditions LC3-II levels in *VPS41*^*S285P/R662\**^ (patient) fibroblasts were similar to control and parental cells (Figure 4A, quantified in 4A’). However, when autophagy was induced by nutrient starvation, LC3-II levels in *VPS41*^*WT/WT*^, *VPS41*^*WT/S285P*^ and *VPS41*^*WT/R662\**^ fibroblasts strongly increased to 150-200% of the original levels, whereas in patient fibroblasts only a mild, 40%, increase was seen. This shows an inability of patient fibroblasts to respond to nutrient starvation via upregulation of autophagy. Thus, although we do not see an increase of LC3 in steady state levels, the patient fibroblasts show a clear defect in regulation of the autophagy response.

**Figure 4.**
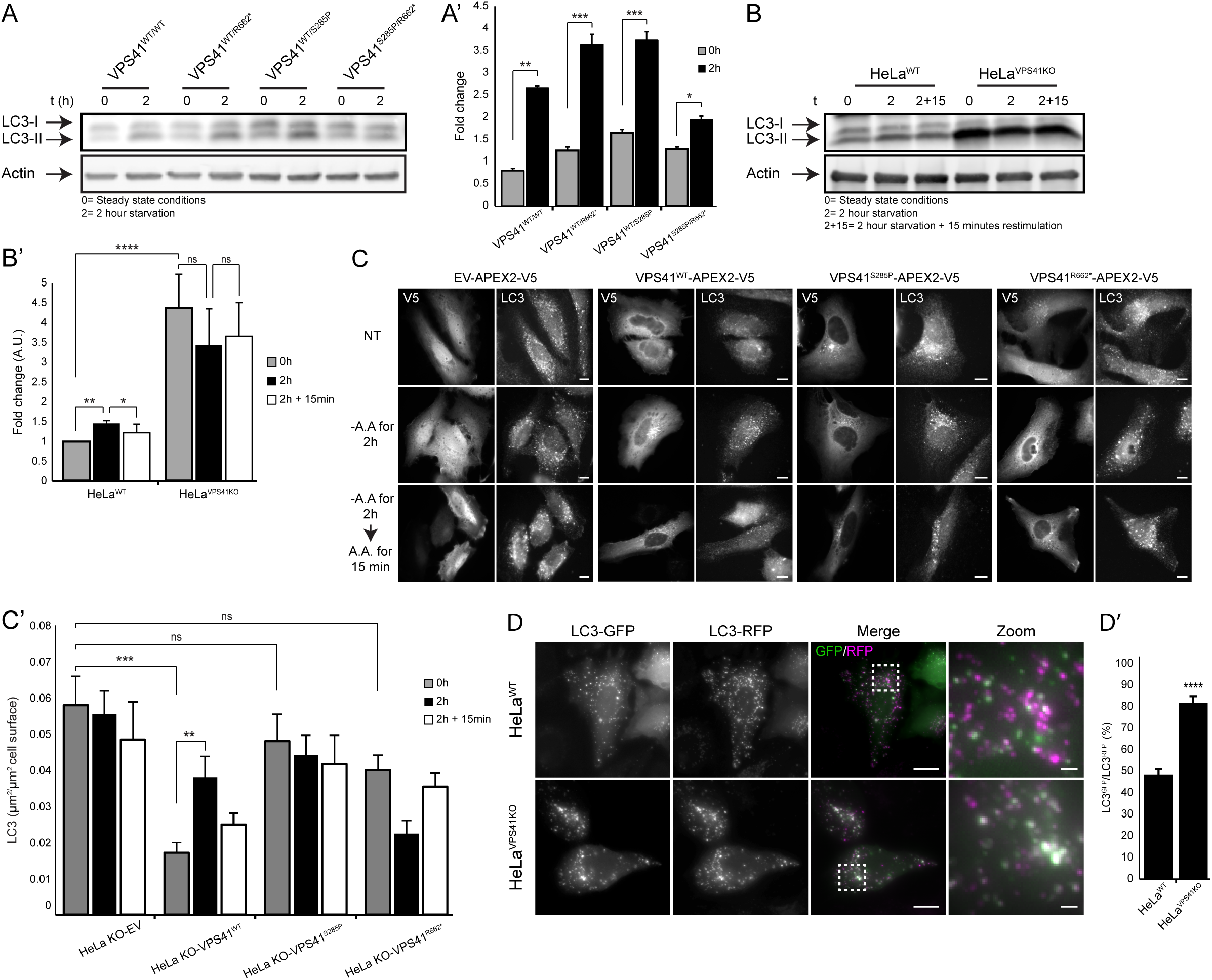
Patient fibroblasts and *VPS41*^*KO*^ cells are impaired in autophagic flux and response to autophagic stimuli. **(A)** Western Blot of LC3 expression levels in control, parental and patient fibroblast. Steady state (0h) levels of lipidated LC3 (LC3-II) are comparable in all conditions. Induction of autophagy by nutrient starvation (2h) results in a 150-200% increase in LC3-II in *VPS41*^*WT/WT*^, *VPS41*^*WT/S285P*^ and *VPS41*^*WT/R662\**^ fibroblasts and only 40% increase in patient fibroblasts (quantified in **A’**) (n=3). Error bars represent the SEM. *p<0.05, **p<0.01, ***p<10^−4^; Unpaired *t* test. **(B)** Western Blot analysis of HeLa^*VPS41KO*^ cells showing a 4-fold increase in LC3-II-protein levels in steady state conditions (0h). In contrast to control cells, nutrient starvation (2h) did not increase LC3-II protein levels in HeLa^*VPS41KO*^ cells indicating irresponsiveness to nutrient availability (quantified in **B’**) (n=3). Error bars represent the SEM. *p<0.05, **p<0.01, ****p<10^−5^; one-ANOVA with Bonferroni correction for multiple comparisons. **(C)** HeLa^*VPS41KO*^ cells transfected with EV-APEX2-V5, VPS41^WT^-APEX2-V5, VPS41^S285P^-APEX2-V5 or VPS41^R662*^-APEX2-V5 and labeled for V5 and LC3 for immunofluorescence microscopy. Rescue with VPS41^WT^-APEX2-V5 decreases the number of LC3-positive compartments in steady state conditions (NT) and restores responsiveness to nutrient starvation (-AA) and replenishment (-AA, +AA). Neither of the mutant VPS41 variants rescues this autophagy phenotype (quantified in **C’**) (n=3). Bars 10µm. Error bars represent the SEM. **p<0.01, ***p<10^−4^; one-ANOVA with Bonferroni correction for multiple comparisons. **(D)** Immunofluorescence of HeLa^*WT*^ and HeLa^*VPS41KO*^ cells transfected with the LC3^GFP/RFP^ tandem construct. A high percentage of the RFP-positive compartments represent autophagosomes (colocalization GFP/RFP) in HeLa^*VPS41KO*^ cells, indicating a block in autophagic flux (quantified in **D’**). Bars 10µm, zoom 1µm. Error bars represent the SEM. Unpaired *t* test, ****p<10^−5^.

To better understand this phenomenon we studied autophagy in HeLa^*VPS41KO*^ cells. When grown under nutrient-rich conditions LC3-II levels in these cells were significantly elevated compared to HeLa^*WT*^ cells (Figure 4B, quantified in 4B’). This is in contrast to our observations in patient fibroblasts, but similar to previous results in VPS41-depleted HeLa cells^54^ and our non-published observations in A549 and PC12 *VPS41*^*KO*^ cells (data not shown). Immunofluorescence on HeLa^*VPS41KO*^ cells also showed more LC3 puncta, indicative for the increase in number of autophagosomes (Figure 4C, quantified in 4C’). We then transfected HeLa^*WT*^ and *VPS41*^*KO*^ cells with an LC3^GFP/RFP^ tandem construct. The GFP and RFP signals co-label autophagosomes, but the GFP signal is lost after autophagosome - lysosome fusion(48). *VPS41*^*KO*^ cells showed a significant higher percentage of GFP/RFP co-localization, indicating that in the absence of VPS41, autophagosome – lysosome fusion is impaired (Figure 4D, quantified in 4D’), resulting in more LC3 puncta. Strikingly, starvation of HeLa^*VPS41KO*^ cells did not further increase LC3-II protein levels or number of LC3 puncta (Figure 4B and 4C), which exactly mimics the unresponsiveness of patient fibroblasts to starvation. Rescue of HeLa^*VPS41KO*^ cells with VPS41^WT^ reduced the number of LC3 puncta and restored the capacity of cells to react to starvation and restimulation (Figure 4C). By contrast, expression of *VPS41*^*S285P*^ or *VPS41*^*R662\**^ did not restore the autophagy phenotype (Figure 4C). This shows that both VPS41 variants are unable to restore basal autophagy levels and provoke an inability to react to nutrient starvation.

We conclude from these data that patient fibroblasts, *VPS41*^*KO*^ cells or *VPS41*^*KO*^ cells expressing *VPS41*^*S285P*^ or *VPS41*^*R662\**^ have multiple defects in autophagy; a decrease in autophagosome - lysosome fusion and a decreased ability to raise LC3-II levels after starvation. Concomitantly, *VPS41*^*KO*^ cells display higher LC3-II levels and contain more autophagosomes under steady state conditions. Patient fibroblasts do not show a markable increase in LC3-II, but this is possibly obscured by the high variation between these primary cultured cells. Alternatively, compensatory mechanisms may have developed. The autophagy defect is specific for the patient derived fibroblasts and not observed in any of the parental cell lines, indicative that this phenomenon is important for disease development.

### VPS41 deficiency causes mTORC1 inhibition

A complex of v-ATPase/Ragulator/Rag GTPases present at the lysosomal membrane senses nutrient status in lysosomes and, in nutrient-rich conditions, recruits the mammalian target of rapamycin complex 1 (mTORC1). Upon nutrient deprivation, mTORC1 dissociates from the lysosomal membrane and loses its kinase activity. Consequently, transcription factors TFEB and TFE3, substrates of mTORC1, translocate to the nucleus where they activate the CLEAR (Coordinated Lysosomal Expression and Regulation) network that induces lysosomal biogenesis and autophagy(49). LC3 is a target of the CLEAR network and mTORC1 inhibition results in enhanced LC3 levels(49). Since patient derived fibroblasts and *VPS41*^*KO*^ cells are less responsive to starvation (Figure 4A-C) this prompted us to investigate mTORC1 activity in *VPS41* patient fibroblasts and KO cells.

We first studied the recruitment of mTORC1 to lysosomal membranes by monitoring colocalization with LAMP-1. Immunofluorescence of *VPS41*^*S285P/R662\**^ patient fibroblasts grown in nutrient-rich conditions showed a striking decrease in localization of mTORC1 to LAMP-1-positive puncta when compared to control or parental cells (Figure 5A and Figure S7, quantified in 5A’). In healthy cells, starvation induces mTORC1 dissociation from lysosomes, whereas restimulation with complete medium restores membrane association. Indeed, in control *VPS41*^*WT/WT*^, paternal *VPS41*^*WT/S285P*^ and maternal *VPS41*^*WT/R662\**^ fibroblasts, mTORC1 dissociated from LAMP-1 positive lysosomes upon 2 hours starvation and partially re-localized after 15 min restimulation. By contrast, mTORC1 localization in *VPS41*^*S285P/R662\**^ patient fibroblasts did not change upon starvation or restimulation (Figure 5A, quantified in 5A’). To test the effect of mTORC1 dissociation on TFEB/TFE3 localization we performed immunofluorescence microscopy of TFE3, which allows for detection of endogenous TFE3. This showed that TFE3 was continuously present in the nucleus of patient cells, regardless of nutrient status. By contrast, *VPS41*^*WT/WT*^, *VPS41*^*WT/S285P*^ and *VPS41*^*WT/R662\**^ fibroblasts only showed nuclear TFE3 localization in response to starvation (Figure 5B, quantified in 5B’). These data indicate that in patient cells mTORC1 is strongly inhibited and TFE3 is continuously activated, which predicts an increase in expression of lysosomal and autophagy proteins(50,51). Western Blot analysis indeed showed that in addition to LC3, protein levels of LAMP-1 and the lysosomal hydrolase cathepsin B were increased in patient’s fibroblasts (Figure S8).

**Figure 5.**
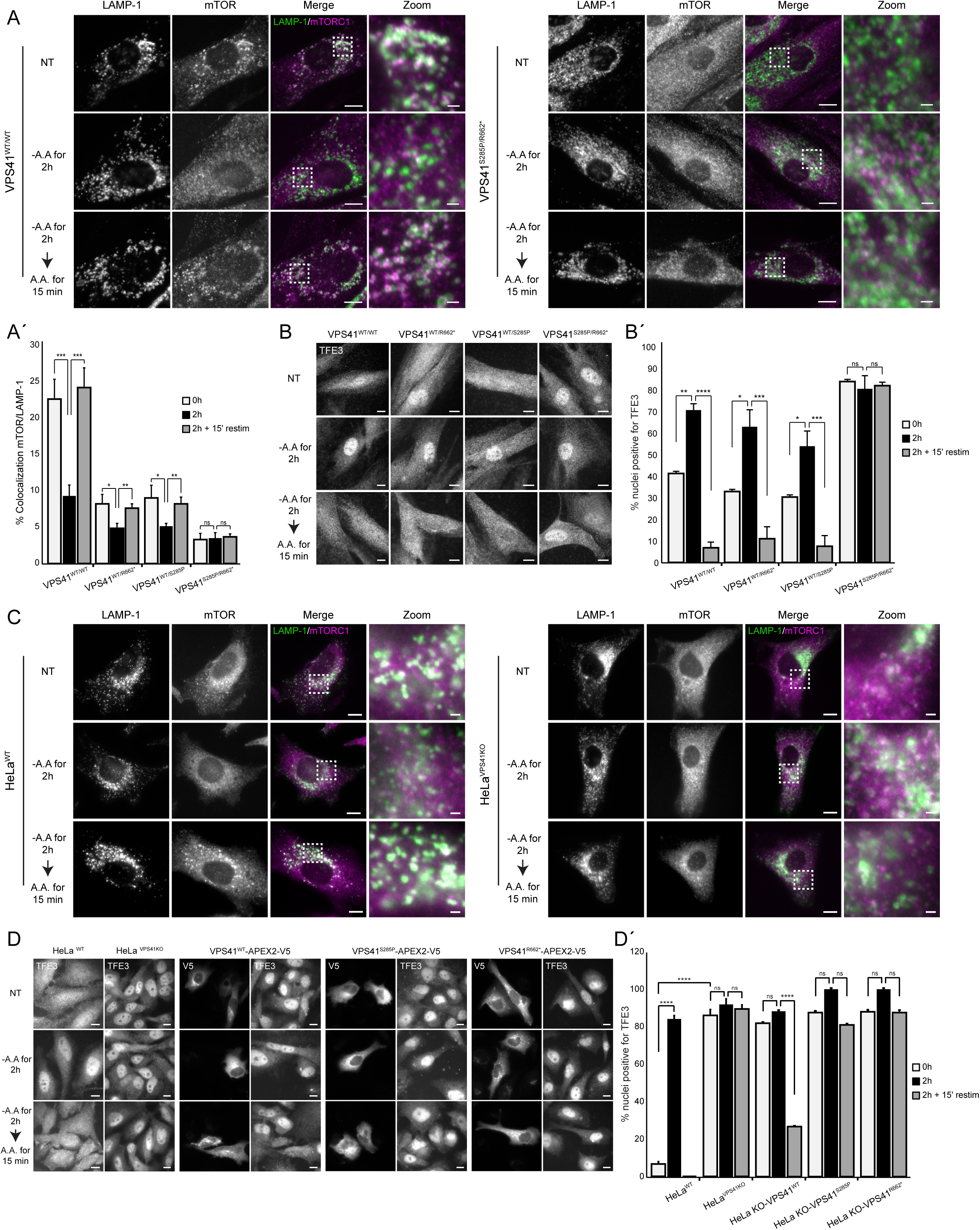
Patient fibroblasts and *VPS41*^*KO*^ cells show mTORC1 inhibition. **(A)** Immunofluorescence of control, parental and patient fibroblasts labeled for LAMP-1 and mTORC1. In steady state conditions (Non Treated (NT)) *VPS41*^*S285P/R662\**^ fibroblasts show less colocalization between mTORC1 and LAMP-1 positive compartments. mTORC1 does not respond to changes in nutrient status. *VPS41*^*WT/WT*^, *VPS41*^*WT/S285P*^ and *VPS41*^*WT/R662\**^ show an appropriate mTORC1 response upon nutrient deprivation (-Amino Acids (-AA)) or stimulation (-AA, +AA) (Figure S6, quantified in **A’**) (n=3). Bars 10µm, zoom 1µm. Error bars represent the SEM. *p<0.05, **p<0.01, ***p<10^−4^; one-ANOVA with Bonferroni correction for multiple comparisons. **(B)** Immunofluorescence of *VPS41*^*WT/WT*^, *VPS41*^*WT/S285P*^, *VPS41*^*WT/R662\**^ and *VPS41*^*S285P/R662\**^ fibroblasts labeled for TFE3. In *VPS41*^*S285P/R662\**^ fibroblasts, TFE3 is constitutively localized in the nucleus, regardless of nutrient state (quantified in **B’**) (n=3) Bars 10µm. Error bars represent the SEM. *p<0.05, **p<0.01, ***p<10^−4^, ****p<10^−5^; one-ANOVA with Bonferroni correction for multiple comparisons. **(C)** HeLa^*WT*^ and HeLa^*VPS41KO*^ cells labeled for LAMP-1 and mTORC1 immunofluorescence. In HeLa^*VPS41KO*^ cells mTORC1 is dissociated from LAMP-1 positive compartments regardless of nutrient state. In HeLa^*WT*^ cells mTORC1 colocalizes with LAMP-1 in steady state and after nutrient restimulation. Bars 10µm, zoom 1µm. **(D)** TFE3 immuno-fluorescence in HeLa^*WT*^ and HeLa^*VPS41KO*^ cells. As in patient fibroblasts, in HeLa^*VPS41KO*^ cells TFE3 is constitutively localized in the nucleus. Rescue with VPS41^WT^-APEX2-V5, VPS41^S285P^-APEX2-V5 or VPS41^R662*^-APEX2-V5. Reintroduction of VPS41^WT^ restores cytoplasmic localization of TFE3 after restimulation with nutrients. Neither of the mutant VPS41 variants could rescue the TFE3 phenotype (quantified in **D’**). (n=3). Bars 10µm. Error bars represent the SEM. ****p<10^−5^; one-ANOVA with Bonferroni correction for multiple comparisons.

When similar experiments were performed on HeLa^*VPS41KO*^ cells we also found a decreased lysosomal localization of mTORC1 (Figure 5C) and continuous nuclear localization of TFE3, independent of nutrient status (Figure 5D). We further refer to this pattern as the mTORC1/TFE3 phenotype. Expression of VPS41^WT^ in HeLa^*VPS41KO*^ cells rescued this phenotype after restimulation, whereas expression of VPS41^S285P^ or VPS41^R662*\**^ had no effect (Figure 5D, quantified in 5D’), inferring that neither of the VPS41 variants could rescue the mTORC1/TFE3 phenotype. These data show that cells lacking VPS41 or expressing VPS41^S285P^ and/or VPS41^R662*\**^, mTORC1 dissociates from lysosomes.

A possible explanation for the mTORC1/TFE3 phenotype is that the defect in HOPS function results in insufficient delivery of nutrients to lysosomes, resulting in nutrient depletion and subsequent mTORC1 dissociation. If so, any block in HOPS function should give a similar phenotype. To test this we made HeLa^KO^ cells for *VPS11* and *VPS18*, which are part of HOPS but not required for the ALP/LAMP pathway(18). Both KO cell lines showed increased LC3-II protein levels in basal conditions and a constitutive nuclear localization of TFE3, independent of nutrient status (Figure S9A and S9B, quantified in S9B’).

We conclude from these data that heterologous expression of VPS41^S285P^ and VPS41^R662*^ or complete KO of *VPS41* results in the dissociation of mTORC1 from lysosomes, a subsequent nuclear localization of TFE3 and increased expression of LC3-II. Moreover, these cells lose the ability to respond to starvation. The mTORC1 inhibition is likely caused by a lack of HOPS function, since depletion of other HOPS subunits result in a similar phenotype.

### VPS41^S285P^ allows for normal regulated secretion in PC12 cells

VPS41 is required for secretory protein sorting and secretory granule biogenesis in a pathway that is independent of the HOPS complex(31). This VPS41 function requires the N-terminal residues 1-36 for interaction with AP-3 and the presence of the C-terminal located CHCR domain(31,52). Since the VPS41^R662*^ variant lacks both the RING domain and part of the CHCR domain, a function in regulated secretion is excluded(31). However, the VPS41^S285P^ mutant could theoretically still exert this HOPS independent function. To test this, we first investigated whether VPS41^S285P^ can interact with AP-3. Pulldowns of recombinant VPS41 constructs with the hinge-ear domain of AP3D1 showed that VPS41^S285P^ binds AP-3 with equivalent affinity as VPS41^WT^ (Figure 6A, S10). To directly test the effect of VPS41^S285P^ on regulated secretion, we performed secretion experiments using *VPS41*^*KO*^ PC12 cells in which we reintroduced either VPS41^WT^ or VPS41^S285P^. Regulated protein secretion was measured as previously described(30). Briefly, cells were incubated in Tyrode’s buffer containing 2.5 mM (basal) or 90 mM (stimulated) K+ and the supernatant and cell lysates were analyzed by quantitative fluorescent immunoblotting. This unequivocally showed that VPS41^S285P^ rescued secretion to the same extent as VPS41^WT^ (Figure 6B-D). As previously showed(31), cellular SgII protein levels were decreased in VPS41^KO^ cells. Reintroduction of VPS41^WT^ or VPS41^S285P^ resulted in an increase in these cellular SgII levels. Together these data show that VPS41^S285P^ binds AP-3 and rescues the regulated secretory pathway in *VPS41*^*KO*^ PC12 cells indicating that the HOPS-independent function of VPS41^S285P^ is not affected.

**Figure 6.**
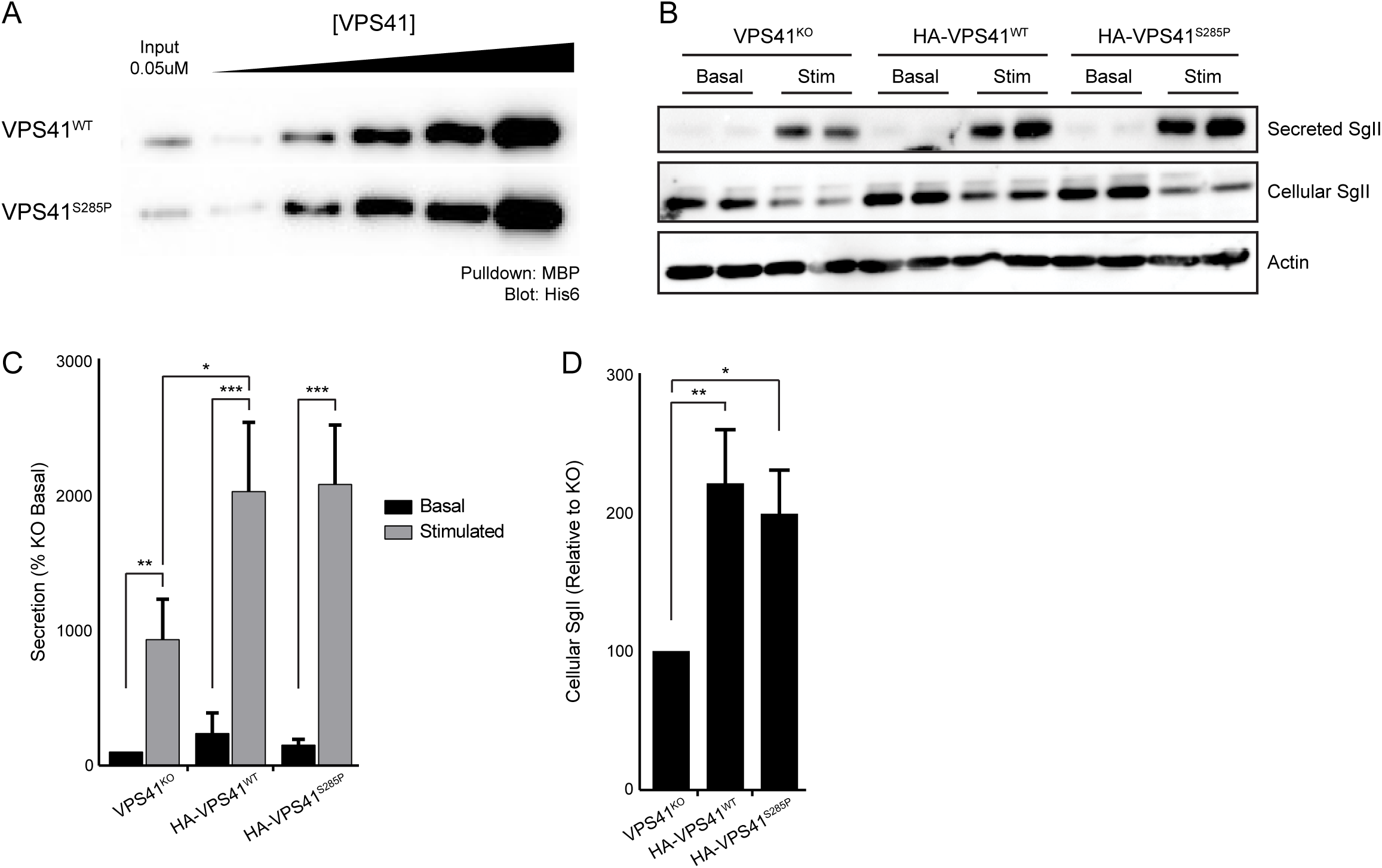
VPS41^285P^ rescues regulated secretion in PC12 VPS41^KO^ cells. **(A)** Western blot of pulldown of AP3 incubated with HIS-6 VPS41^WT^ or VPS41^S285P^. There is no difference in affinity between VPS41^WT^ and VPS41^S285P^ (Figure S9). **(B)** PC12 cells VPS41^KO^, or VPS41^KO^ transduced with HA-VPS41^WT^ or HA-VPS41^S285P^ lentivirus were washed and incubated for 30 min in Tyrode’s solution containing 2.5 mM K+ (basal) or 90 mM K+ (stimulated). Cellular **(C)** and secreted **(D)** secretogranin II (SgII) were measured by quantitative fluorescence immunoblotting. Error bars represent the SEM. (n=3). *p<0.05, **p<0.01, ***p<0.001; secretion data analyzed by one-way ANOVA.

### Co-expression of VPS41^S285^ and VPS41^R662*^ abolishes neuroprotective function in a *C. elegans* model of Parkinson’s disease

The clinical symptoms of the two *VPS41* patients (Figure 1) overlap with PD(53,54). Previously, a screen for neuroprotective factors in a transgenic *C. elegans* model for PD showed that overexpression of human *VPS41* protects against α-synuclein induced neurodegeneration(32–34). The neuroprotective effect of VPS41 depends on the presence of the WD40 and CHCR domains and ability to interact with Rab7 and AP-3(32,35). To explore the impact of the disease-causing VPS41^S285P^ and VPS41^R662*^ mutants in this PD model, we made transgenic nematodes co-expressing pairs of human VPS41 isoforms (hVPS41) in dopaminergic neurons and crossed these with isogenic strains overexpressing either GFP alone or together with human α-synuclein to mimic PD.

First we combined VPS41^WT^ with VPS41^S285P^ or VPS41^R662^ expression, reflecting paternal and maternal conditions, and studied the effect on neurodegeneration in the absence of α-synuclein. At 7 or 10 days post-hatching, there was no statistically significant change in neurodegeneration in any of these strains (Figure 7A,B). Likewise, when we co-expressed VPS41^S285P^ with VPS41^R662*^ there was no discernable increase in neurodegeneration. This demonstrates that co-expression of the two patient-specific hVPS41 variants by itself does not increase neurodegeneration in (ageing) animals.

**Figure 7.**
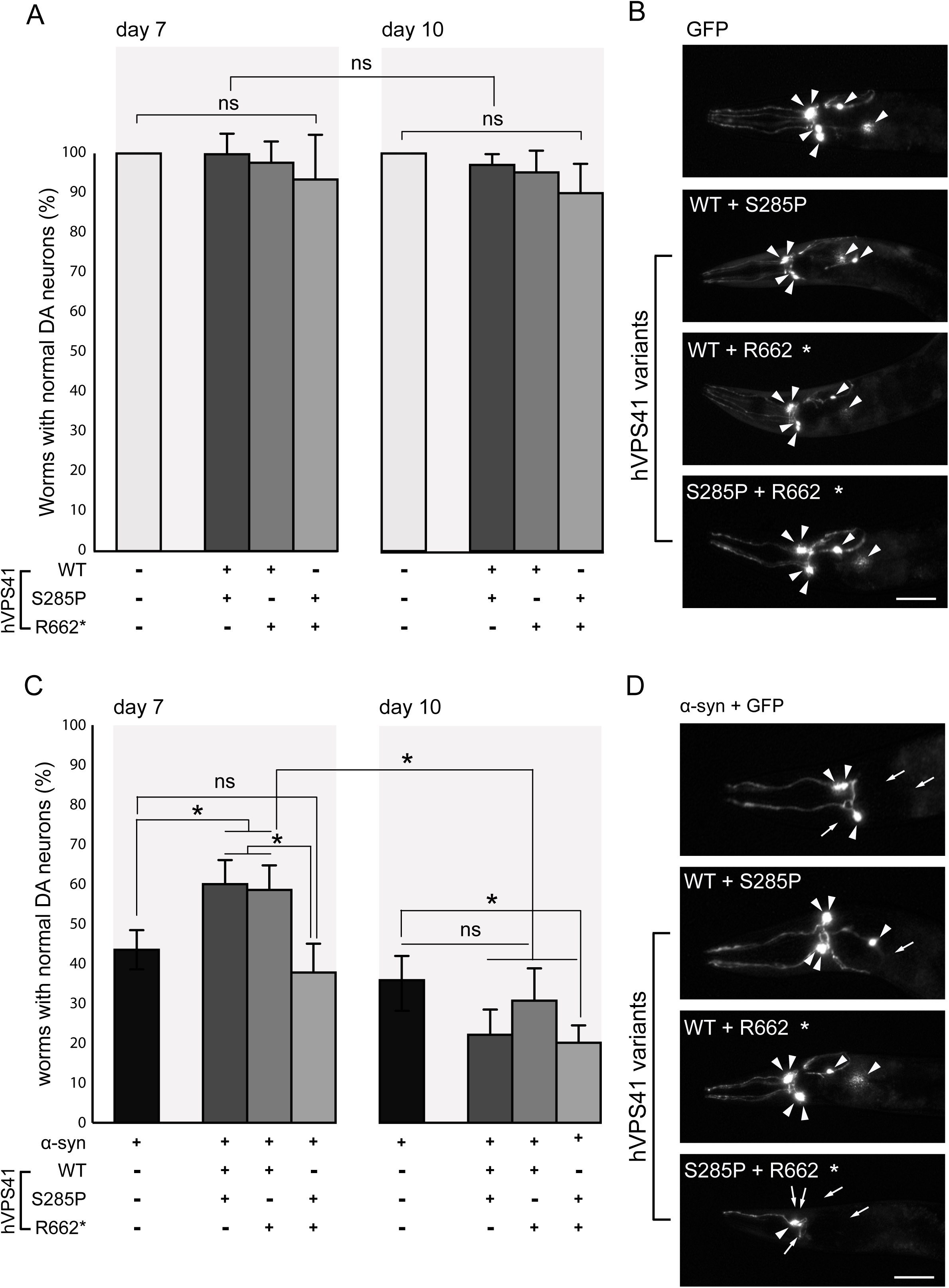
Compound heterozygous expression of hVps41 variants fails to rescue α-synuclein-induced neurodegeneration. **(A)** Graph representing the percentage of animals with 6 normal dopaminergic neurons. Heterozygous expression with hVPS41^WT^ (strains UA386 or UA387) or compound heterozygous expression of hVPS41 variants (strain UA388) does not yield significant changes in neurodegeneration compared to the P_*dat-1*_::GFP (strain BY250) control at either day 7 or day 10 post-hatching. Error bars indicate standard deviation. n=30 adult worms for each of 3 independent experiments for GFP (total of 90 worms) and n=90 for each independent transgenic strain (30 worms/trial x 3 independent transgenic lines = 270 worms) for each of 3 independent experiments. **(B)** Representative images of the neuroprotection assay described in part A, where worms express GFP specifically in the anterior 6 DA neurons. Intact DA neurons are indicated with arrowheads. hVPS41 variant backgrounds are expressed as indicated. Magnification bar = 20 μm. **(C)** Graph representing the percentage of animals with 6 normal dopaminergic neurons in the anterior region of P_*dat-1*_::GFP; P_*dat-1*_::α-syn (strain UA44) animals with heterozygous expression of hVPS41 variants. Heterozygous expression of hVPS41^WT^ with either variant (strains UA389 or UA390) significantly rescues neurons from α-syn-induced degeneration at day 7 whereas compound heterozygous expression (strain UA391) fails to rescue neurodegeneration (*p<0.05, One-Way ANOVA with Tukey correction for multiple comparison). At day 10, none of the heterozygous backgrounds significantly rescue α-synuclein-induced neurodegeneration. Error bars indicate standard deviation. n=30 adult worms for each of three independent experiments for the α-synuclein (α-syn) strain (total of 90 worms) and n=90 for each independent transgenic strain (30 worms/trial x 3 independent transgenic lines = 270 worms) for each of 3 independent experiments. **(D)** Representative images of DA neurons from *C. elegans* expressing P_*dat-1*_::GFP; P_*dat-1*_::α-syn, with or without hVPS41 variants, as described in part C. Intact DA neurons are indicated with arrowheads and missing neurons are indicated with arrows. Magnification bar = 20 μm.

We then analyzed VPS41 neuroprotective function in the strains overexpressing α-synuclein. At 7 days post-hatching, only ∼43% of the animal population overexpressing α-synuclein retained the full complement of dopaminergic neurons. By 10 days, this had decreased to only ∼33% of the population. Co-expression of VPS41^WT^ with either VPS41^S285P^ or VPS41^R662*^ significantly enhanced the neuroprotection at 7 days post-hatching, but at day 10 the protective effect was lost (Figure 7C,D).

By contrast, animals co-expressing VPS41^S285P^ and VPS41^R662*^, reflecting patient conditions, showed no protective effect at all at either day 7 or 10 (Figure 7C,D). Therefore, the protective effect observed with the compound heterozygous combinations is lost as animals age. By contrast, expression of VPS41^WT^ alone rescues neurodegeneration also at day 10, albeit it with reduced efficiency compared to day 7(32,34). Moreover, neuroprotection at day 10 was significantly diminished in worms co-expressing both of the clinical VPS41 variants in combination indicating additional stress besides α-synuclein overexpressing and aging alone. These data collectively indicate that the neuroprotective capacity of VPS41 is differentially impacted by genetic composition.

## Discussion

The vacuolar protein sorting-associated protein 41 (VPS41) has a major contribution in the vesicle-mediated trafficking to lysosomal compartments including endocytic transport and the autophagic pathway. VPS41 is part of the HOPS complex, a multisubunit tethering complex which mediates fusion of lysosomes with late endosome and autophagosomes(12,13). Independently of the HOPS complex, VPS41 is also involved in the transport of lysosomal membrane proteins from the Trans-Golgi network independently of HOPS(18), and plays a role in regulated secretion of neuropeptides(31). VPS41 is previously identified as neuroprotector in *C. elegans* and mammalian models of Parkinson’s Disease (PD)(32,34,35). However, to our best knowledge, no specific human condition has been reported in association with a pathogenic variant in VPS41. We here report on two siblings with compound heterozygote variants in VPS41 and show the implications of these variants on a cellular level. We show that both VPS41 variants result in a nonfunctional HOPS complex, causing a defect in the delivery of endocytic and autophagy cargo to enzymatically active lysosomes. Most strikingly, VPS41 dysfunction causes dissociation of the mTORC1 complex from lysosomes which renders cells unable to respond to nutrient starvation and results in increased autophagy levels. Our studies thereby for the first time link HOPS complex functionality to mTORC1 signaling.

The patients, two brothers, were diagnosed with global developmental delay, hypotonia, ataxia and dystonia. By exome sequencing we identified a missense mutation in the WD40 domain (VPS41^S285P^), and a nonsense mutation in the CHCR domain resulting in a premature stopcodon (VPS41^R662*^). Neither VPS41 mutant forms a functional HOPS complex, indicating that the patient cells lack HOPS functionality. The R662* variant lost binding affinity to HOPS components VPS18 or VPS33A due to a lack of the RING domain(42), yet was recruited to late endosomes and lysosomes by interaction with Rab7 and Arl8b. These data imply that recruitment of VPS41 to endo-lysosomal membranes does not require HOPS complex incorporation. However, VPS41^R662*^ is not expressed in patient cells, indicating that the disease phenotype is caused by the VPS41^S285P^ mutation solely. VPS41^S285P^ interacted with other HOPS complex subunits and was recruited to LAMP-1 positive endo-lysosomal membranes, but did not rescue the fusion defect in *VPS41*^*KO*^ HeLa cells. Since a substitution of serine to proline is predicted to induce a protein folding defect, the most likely explanation is that VPS41^S285P^ has a structural defect that prevents formation of a functional HOPS complex.

A most striking finding in our studies is that patient-derived fibroblasts and *VPS41*^*KO*^ HeLa cells show a decreased lysosomal localization of mTORC1 and a constitutive nuclear localization of TFE3. Accordingly, *VPS41*^*KO*^ cells and *VPS41*^*S285P/R662\**^ fibroblasts did not respond to starvation or subsequent replenishment of nutrients. Neither VPS41 mutant could rescue this mTORC1/TFE3 phenotype in *VPS41*^*KO*^ cells. Defects in mTORC1 signaling have been reported in many neurodegenerative disorders, including Alzheimer and PD(55). However, in contrast to the VPS41 phenotype these conditions show a decrease in lysosomal performance and an increase in mTORC1 activity. This is why development of therapeutics mostly focus on mTORC1 inhibition or TFEB/TFE3 activation(56,57). Our data show that lysosomes in *VPS41* patient fibroblasts or KO cells are acidified and enzymatically active, thus in principle are functionally equipped. However, the transfer of endocytosed and autophagic cargo to these active lysosomes is compromised. A possible explanation for the mTORC1/TFE3 phenotype is therefore that lysosomes sense that they are deprived of nutrients even in nutrient-rich conditions. This ‘starvation’ scenario predicts that other factors blocking late endosome - lysosome fusion also lead to the mTORC1/TFE3 phenotype. Indeed, TFE3 translocates to the nucleus when cells are treated with chloroquine(50), which recently was shown to cause a block in late endosome - lysosome fusion(58). Also inhibition of lysosomal proteases and thereby suppressing degradation of the lysosomal content results in mTORC1 inhibition and autophagy initiation(56,59). Moreover, we found that knockout of the HOPS components *VPS11* or *VPS18*, also leads to TFE3 translocation and increased LC3-II levels, independent of nutrient status, whereas previously a link between VPS39 and mTORC1 signaling was reported(60). These data indicate that a defect in HOPS function generally leads to mTORC1 inhibition.

The mTORC1 dissociation observed in *VPS41* patient and KO cells induced a continuous nuclear localization of TFE3. TFE3 is part of the MiTF/TFE family of transcription factors that also comprises TFEB(61). Similarly to TFEB, TFE3 translocates to the nucleus upon nutrient starvation and binds CLEAR elements in promotors of autophagic and lysosomal genes(49,62). Previous studies had already shown that depletion of *VPS41* decreases HOPS-dependent fusion between autophagosomes and endo-lysosomes, resulting in accumulation of LC3-positive autophagosomes(63). We here recapitulated this finding in HeLa^*VPS41KO*^ cells. Moreover, in HeLa, A549 and PC12 *VPS41*^*KO*^ cells we found by EM an increase in number of autophagosomes (data not shown). Together these data show that depletion of *VPS41* increases LC3 levels by increased protein synthesis as well as decreased lysosomal clearance. In contrast to *VPS41*^*KO*^ cells, LC3-II protein levels in patient-derived fibroblasts were not enhanced. Possibly, the high variability between primary cells obscures changes in LC3-II levels. Alternatively, patient cells may have developed compensatory protective mechanisms against a block in clearance of autophagosomes.

By immuno-EM we found no effect on the concentration of LAMP-1 or LAMP-2 in lysosomal membranes. Also, cathepsin B and cathepsin D were readily activated, marking their presence in the lysosomes. By contrast, a short pulse of Dextran (2 hours) showed a decrease in delivery of endocytosed cargo to enzymatically active lysosomes. Since resident lysosomal proteins have half-lives of several days, this suggests that the absence of HOPS causes a delay rather than a block in cargo delivery. Indeed, prolonged incubation times (72 hours) showed similar co-localization levels of Dextran with SiR-Lysosome cathepsin D in *VPS41*^*KO*^ compared to *VPS41*^*WT*^ cells (data not shown). Of note, since the continuous activation of TFE3 leads to an increase in biosynthesis of lysosomal proteins, we cannot rule out that potential defects in the kinetics or efficiency in transport of LAMPs and other lysosomal membrane proteins are compensated for by a higher production. Future studies using pulse-chase live cell imaging and mass spectrometry on isolated lysosomes are required to address the kinetics of lysosomal membrane protein transport in *VPS41*^*KO*^ cells. Concurrent with a lack of HOPS function we found that transfer of endocytosed cargo from late endosomes to lysosomes is compromised in patient-derived fibroblasts. In the paternal *VPS41*^*WT/S285P*^ fibroblasts there was no endocytosis defect, indicating that the WT allele compensates for VPS41^S285P^ function. In the maternal *VPS41*^*WT/R662\**^ fibroblasts, we did find a defect in transfer of endocytosed cargo to lysosomes, but this was not accompanied by the mTORC1/TFE3 phenotype. Cells were still able to respond to nutrient starvation and re-stimulation, indicating that the overall defect in lysosomal fusion is less severe than in patient cells and that the combined input of endocytic and autophagy cargo into lysosomes is sufficient to activate the mTORC1 complex.

Independent of HOPS, VPS41 is required for formation of secretory granules as well as transport of lysosomal membrane proteins via the ALP/LAMP-carrier pathway(18,19,23,64). Both functions depend on the interaction of VPS41 with AP-3(20–24). By co-IP we found that VPS41^S285P^ still binds AP-3. In addition, VPS41^S285P^ was able to rescue the secretion phenotype when expressed in *VPS41*^*KO*^ PC12 cells. These data indicate that HOPS-independent functions of VPS41 are not significantly affected in patient cells due to the expression of the paternal *VPS41*^*S285P*^ allele. Notably, since *VPS41* knockout mice are embryonic lethal(14), the HOPS-independent functions of VPS41^S285P^ may be a vital factor for the survival of the patients.

VPS41 was previously reported to confer neuroprotection in both C. elegans and mammalian models for PD(32–35). Overexpression of α-synuclein results in degeneration of dopaminergic neurons, which is prevented by simultaneous overexpression of human VPS41. Both the WD40 and CHCR domain of VPS41 are necessary for this neuroprotection(32,34), which are mutated in VPS41^S285P^ and VPS41^R662*^, respectively. We found that transgenic nematodes co-expressing hVPS41^WT^ with either VPS41^S285P^ or VPS41^R662*^retained the neuroprotective effect, resulting in a decrease in neurodegeneration. However, co-expressing VPS41^S285P^ and VPS41^R662*^ did not result in neuroprotection and even exacerbated cell death. Since we found that the VPS41 patient cells completely lack HOPS function, the neurodegenerative phenotype in the nematodes is likely HOPS related. Moreover, since HOPS-dependent autophagosome-lysosome fusion is important for clearance of aggregated material, which particularly tends to accumulate in neurons over the course of ageing(17,43,45,47), overexpression of VPS41 in the *C. elegans* model possibly results in increased clearance of aggregated or misfolded proteins through autophagy. The intersection of lysosomal dysfunction and neurodegeneration has become a critical junction in the mechanistic underpinnings of PD(65). This is best exemplified by the relationship between homo- vs. heterozygous inheritance of mutations in the Gaucher’s disease associated gene, GBA1, which has become the most prevalent genetic risk factor for PD(66). In the context of these results, it is tempting to speculate that an increased understanding of autolysosomal trafficking defects in VPS41 patient cells may serve to inform therapeutic strategies for neurodegenerative disorders as well(35).

Currently, patients with mutations in HOPS core components *VPS11, VPS16* and *VPS33A* have been reported in literature, which all show a neurodegenerative phenotype(11,67–74). Moreover, during our studies, a third patient with compound heterozygote mutations in *VPS41* was identified (Personal communication by I. Stolte-Dijkstra, University Medical Center Groningen), bearing the VPS41^R662*^ mutation as well as a missense mutation (VPS41^H466R^), and displaying a similar neurological phenotype. By western blot the VPS41 protein levels were markedly decreased in the patient-derived fibroblasts and similar TFE3 nuclear translocation was observed (data not shown), indicating that this patient also suffers from a loss of HOPS function.

In summary, our data suggest that the *VPS41* patients may represent a novel class of lysosomal disorders in which lysosomes are enzymatically active but, due to a trafficking defect, insufficiently reached by nutrients, causing continuous ‘starved’ lysosomes. Interestingly, this results in an inhibition of mTORC1 as opposed to the hyperactivity of mTORC1 seen in many other neurodegenerative diseases.(75–77). These observations are of particular importance with regards to the design of a possible treatment of HOPS related disorders(69,70,73,78,79).

## Methods and Materials

### Identification of *VPS41* variants

Whole exome sequencing was performed on both affected siblings and their father. Briefly, total leukocyte DNA was enriched using the Agilent SureSelect Human All Exon 50 Mb capture kit (Agilent Technologies, SantaClara, CA). Bidirectional 100-bp paired-end sequencing (Illumina Hi-Seq 2000; Illumina, Inc., San Diego, CA) was performed. Sequence reads were aligned to the hg19 reference human genome using the Burrows-Wheeler Aligner (http://bio-bwa.sourceforge.net), and duplicate reads were removed with MarkDuplicates (Picard toolsversion 1.35; http://picard.sourceforge.net). Alignments were refined using local realignment in Short Read Micro re-Aligner version 0.1.15(http://sourceforge.net/projects/srma). Single nucleotide variants (SNVs) and insertion/deletions (indels) were detected using Genome Analysis Toolkit version 1.1.28 (http://www.broadinstitute.org/gatk). SNVs and indels were annotated with SnpEff (http://snpeff.sourceforge.net/) and filtered to remove common (frequency.0.01) polymorphisms by screening against the following databases: dbSNP (http://www.ncbi.nlm.nih.gov/projects/SNP/), 1000 Genomes (www.1000genomes.org), the NHLBI Exome Variant Server (http://evs.gs.washington.edu/EVS/), and data from 130 local control exomes. Filtered variants were annotated with SIFT (http://sift-dna.org) and PolyPhen-2 (http://genetics.bwh.harvard.edu/pph2/). Variants were confirmed by Sanger sequencing according to standard protocols in both siblings, their father and their mother.

### Antibodies and Reagents

Antibodies used in this study: mouse anti-LAMP-1 CD107a (555798, BD Pharmingen), rabbit anti-LC3 (NB600-1384, Novus Biologicals), rabbit anti-p70 S6 Kinase and Phospho-p70 S6 Kinase (Thr389) (9202 and 9205 respectively, Cell Signaling), goat anti-cathepsin B (AF953, RD Systems), goat anti-cathepsin D (AF1014, RD systems), mouse anti-VPS41 (SC-377271, Santa Cruz), mouse anti-Actin (69100, MP Biomedicals), mouse anti-GFP (11814460001, Roche), rabbit anti-GFP (A6455, Invitrogen), rabbit anti-FLAG (F7429, Sigma), rabbit anti-V5 (V8137, Sigma), mouse anti-V5 (R960-25, Invitrogen), rabbit anti-mTOR (2983 Cell Signaling), rabbit anti-TFE3 (14779, Cell Signaling), mouse anti-Hsp65 and rabbit anti-VPS18 (ab178416, Abcam). Secondary antibodies used are: goat anti-mouse-ALEXA568 (A11031, ThermoFisher), goat anti-mouse-ALEXA488 (A11029, ThermoFisher), goat anti-rabbit-ALEXA568 (A11036, ThermoFisher), donkey anti-rabbit-ALEXA488 (A21206, ThermoFisher), IRDye^®^ 800CW goat anti-rabbit IgG (925-32211, LI-COR), IRDye^®^ 800CW goat anti-mouse IgG (925-32210, LI-COR), IRDye^®^ 680RD goat anti-rabbit IgG (925-68071, LI-COR), goat anti-mouse IgG-ALEXA680 (A28183, ThermoFisher). As bridging antibody between mouse antibodies and protein-A gold, we used rabbit anti-mouse IgG (610-4120, Rockland). Antibodies were visualized for EM using protein-A gold conjugated to 10 or 15nm gold particles (Cell Microscopy Core, UMC Utrecht).

### Endocytosis and lysosomal functionality assays

To study endocytosis, cells were incubated for 2h with 10.000MW Dextran-ALEXA Fluor 568 (Invitrogen) in a 1:100 dilution. Cells were washed 5 times using warm PBS, fixed with 4% PFA, embedded in Prolong DAPI (Invitrogen) and imaged. Lysosomal acidity was determined using Lysotracker™ Red (Molecular Probes) in a 1:100 for 30min after which cells were washed 5 times using warm PBS, fixed with 4% PFA, embedded in Prolong DAPI (Invitrogen) and imaged. To visualize active lysosomes, we used the SiR-lysosome (Spirochrome) probe in a 1:1000 concentration. Cells were incubated for 3h, washed with PBS, fixed with 4% PFA, embedded in Prolong DAPI (Invitrogen) and imaged. For the visualization of Cathepsin B active compartments MagicRed cathepsin B substrate (Molecular Probes) was using in a 1:260 dilution. Cells were incubated for 30min, washed 5 times using warm PBS, fixed with 4% PFA, embedded in Prolong DAPI (Invitrogen) and imaged.

### Cell culture and light and electron microscopy

Patient derived fibroblasts and HeLa cells were cultured in High Glucose Dulbecco’s Modified Eagle’s Medium (Invitrogen) supplemented with 10% FCS in a 5% CO_2_-humidified incubator at 37°C. Cells were transfected using X-tremeGENE™ (Roche) according to manufacturer’s instructions. To induce autophagy, cell were starved for 2h at 37°C using EBSS (ThermoFisher) and restimulated for 15min with complete medium.

Cells destined for immunofluorescence were washed with PBS once and fixed using 4% wt/vol paraformaldehyde (PFA, Polysciences Inc.) for 20min. Cells were washed three times with PBS and permeabilized with 0.1% Triton X-100 (Sigma) for 5min. Samples were blocked for 15min using a 1% BSA solution. Samples were labeled, embedded in Prolong DAPI (Invitrogen) and imaged. When imaging transfected cells, cells with similar transfection levels were selected. Images were taken using a Deltavision wide field microscope (Applied Precision) with a 100/1.4A immersion objective. Image analysis was performed using Volocity software (Perkin Elmer) and macros written in Fiji(80).

For plastic EM, cells grown in 6cm dishes were fixed in 2% wt/vol PFA, 2.5% wt/vol GA in Na-cacodylate buffer (Karnovsky fixative) for 2h at RT. Subsequently, the fixative was replaced for 0.1M Na-cacodylate buffer, pH 7.4. Post fixation was performed using 1% wt/vol OsO4, 1.5% wt/vol K3Fe(III)(CN)6 in 0.065M Na-cacodylate buffer for 2h at 4°C. Next, cells were stained with 0.5% uranyl acetate for 1h at 4°C, dehydrated with ethanol and embedded in Epon. Ultrathin sections were stained with uranyl acetate and lead citrate. Images were taken on a TECNAI T12 electron microscope. For immuno-EM, cells were grown on 6cm dishes. Cells were incubated with BSA-Au^5^ for 2h in culture medium at 37°C, washed using medium and fixed by adding freshly made 2% FA, 0.2% GA in 0.1M phosphate buffer (pH 7.4) by adding an equal amount of fixative to the medium for 5 min. Cells were postfixed using 2% FA, 0.2% GA in 0.1M phosphate buffer which was left on overnight. Cells were washed with PBS/0.05M glycine, scraped using 1% gelatin/PBS, pelleted in 12% gelatin/PBS, solidified and cut into small blocks. Blocks were infiltrated in 2.3M sucrose overnight at 4°C, mounted on aluminum pins and frozen in liquid nitrogen(81). To pick up ultrathin sections, a 1:1 mixture of 2.3M and 1.8% methylcellulose was used(82).

### CRISPR/Cas9 HeLa^*VPS41KO*^

Hela cells were transiently transfected with pSpCas9(BB)-2A-GFP (PX458)(83) encoding sgRNAs targeting VPS41 using X-tremeGENE (Merck) according to the manufacturers recommendation. sgRNAs were designed with http://crispor.tefor.net/(84) and target coding sequences in the N-terminal region encoded by the third coding exon. The sgRNA used is sgRNA2 AAGTATTTCAGTTACCCCAT. GFP-positive cells were sorted using a FACSAria II flow cytometer (BD) and plated in 10 cm dishes. Colonies were picked from these plates after 1 week and expanded. To confirm VPS41 absence, total cell lysates were analyzed by Western blotting using mouse anti-VPS41 (SC-377271, Santa Cruz).

### Co-Immunoprecipitation and Western blotting

Cells were seeded in 15cm dishes and transfected with the appropriate constructs as indicated in the respective figures and according to above mentioned transfection protocols. Cells were washed three times with ice-cold PBS and lysed using a CHAPS lysis buffer (50 mM Tris pH=7.5, 150mM NaCl, 5mM MgCl2, 1mM DTT, 1% (w/v) CHAPS) complemented with protease and phosphatase inhibitor (Roche). Cells were scraped, collected and spun down at 13000rpm for 15 min at 4°C. Protein levels were equalized between samples using a Bradford Protein Assay (Bio-Rad). 10% of the sample was saved as input control. The remainder of the samples was incubated for 1hr with uncoated protein G beads (Millipore) to remove a-specifically bound proteins. Beads were spun down and the supernatant was incubated with beads together with 2µg antibody against the prey. Samples were incubated overnight, extensively washed and eluted using SDS sample buffer and run on precast gradient (4-20%) gels (Bio-Rad). Gels were transferred using Trans-Blot^®^ Turbo™ RTA Mini PVDF Transfer Kit and the Trans-Blot^®^ Turbo™ Transfer system (Bio-Rad). Membranes were blocked with Odyssey^®^ Blocking Buffer (LI-COR) in PBS for 1h at RT and incubated with primary antibody diluted in Odyssey^®^ Blocking Buffer (LI-COR) in 0.1% TBST overnight at 4°C, rocking. Membranes were washed extensively with 0.1% TBST and incubated with secondary antibodies, diluted in Odyssey^®^ Blocking Buffer (LI-COR) in 0.1% TBST at RT for 1h, rocking. The membranes were again washed extensively with 0.1% TBST, followed by 2 washing steps with PBS and 1 washing step with MiliQ. Membranes were scanned using the Amersham™ Typhoon™ Laser Scanner (GE Healthcare Life Sciences).

### Plasmids

The GFP-VPS41 constructs were cloned from hVPS41 cDNA (Origene) into a pDonor201 vector (Invitrogen) using PCR. Next, using the Gateway system and GFP-pcDNA-Dest53 (kindly provided by Prof. R. Roepman, Radboud University Medical Center Nijmegen), a recombination reaction was performed. The VPS41 mutant variants were made using the QuickChange Site-Directed Mutagenesis kit (Agilent) using the primers 5’-CAGCTTGTTGTACTTCCGTATGTAAAGGAGA and 5’- TCTCCTTTACATACGGAAGTACAACAAGCTG to generate GFP-VPS41^S285P^ and 5’- GTTTATCTTCTGAGCTGAATGGGTTAATAGCC and 5’-GGCTATTACCCATTCAGCTCAGAAGATAAAC to generate GFP-VPS41^R662*^. The pcDNA3-APEX2-V5 construct (kindly provided by Dr. S. de Poot) was used as empty vector in pulldown experiments. VPS41-APEX2-V5 was generated via a BamHI insert using pEGFP-C1-mCherry-VPS41 (kindly provided by Prof. J.J.C. Neefjes, University Medical Center Leiden) using the primers 5’-AGAGGGATCCATGGCGGAAGCAGAGGAGCAG and 5’- AGAGTCTAGACTATTTTTTCATCTCCAAAATTG. VPS41^S285P^-APEX2-V5 and VPS41^R662*^-APEX2-V5 were generated via an identical approach using pEGFP-C1-mCherry-VPS41^S285P^ and pEGFP-C1-mCherry-VPS41^R662*^. These VPS41 variants were made using the above mentioned Mutagenesis kit (Agilent) and the same primer pairs.

For the *C. elegans* experiments, the wild-type (wt) P_*dat-1*_::hVPS41 construct was assembled as previously described^32^. Briefly, human VPS-41 cDNA was obtained from Open Biosystems (Huntsville, AL). Plasmid entry vectors were generated using Gateway Technology (Invitrogen) to clone PCR amplified constructs into pDONR221 to generate a hVPS41 entry clone. The hVPS41 entry clone was used to clone hVPS41 into the Gateway expression vector, pDEST-P_*dat-1*_^42^. The mutant hVPS41 alleles were generated by site-directed mutagenesis of the P_*dat-1*_::hVPS41 construct using the primers 5’- ATTCTACATCAGTGGACTTGCACCTCTCTGTGATCAGCTTGTTGTACTTCCGTATGTAAAGGAGATTTCAGAAAAAACGGAAAGAGAATACTGTGCCAG and 5’- CTGGCACAGTATTCTCTTTCCGTTTTTTCTGAAATCTCCTTTACATACGGAAGTACAACAAGCTGATCACAGAGAGGTGCAAGTCCACTGATGTAGAAT to generate the P_*dat- 1*_::hVPS41^S285P^ construct and 5’- GATCTGTCAACAGAGAAACTTTGTAGAAGAGACAGTTTATCTTCTGAGCTGAATGGGTAATAGCCGAAGTGCCCTGAAGATGATTATGGAGGAATTACA and 5’-TGTAATTCCTCCATAATCATCTTCAGGGCACTTCGGCTATTACCCATTCAGCTCAGAAATAAACTGTCTCTTCTACAAAGTTTCTCTGTTA CAGATC to generate the P_dat-1_::hVPS41^R662*^ construct. The identities of the Gateway entry and expression constructs were validated by DNA sequencing.

### *C. elegans* strains

Transgenic *C. elegans* strains were generated by directly injecting a P_*unc-54*_::tdTomato co-injection marker (50ng/µL) and P_*dat-1*_::hVPS41 wt (25ng/µL) with P_*dat-1*_::hVPS41^S285P^ (25ng/µL) or P_*dat- 1*_::hVPS41^R662*^ (25ng/µL) constructs or P_*dat-1*_::hVPS41^S285P^ (25ng/µL) with P_*dat-1*_::hVPS41^R662*^ (25ng/µL) into the gonads of strain N2 Bristol hermaphrodites. Progeny of injected animals were screened for expression of the hVPS41 expression constructs by evidence of fluorescence in the body-wall muscles from the P_*unc-54*_::tdTomato co-expression marker. Several stable lines of each were isolated and crossed with isogenic UA44 (*baIn11*[P_*dat-1*_::α-syn, P_*dat-1*_::GFP]) and BY250 (*vtIs7*[P_*dat-1*_::GFP]) males. Heterozygous hermaphroditic progeny expressing both tdTomato and GFP were transferred and allowed to self-fertilize. Several lines of each P_*dat-1*_::hVPS41wt + P_*dat-1*_::hVPS41^S285P^ [baEx215] in UA44 (UA389) and BY250 (UA386) were analyzed for neurodegeneration at days 7 and 10. Likewise, P_*dat- 1*_::hVPS41wt + P_*dat-1*_::hVPS41^R662*^[baEx216] in UA44 (UA390) and BY250 (UA387) was analyzed. Additionally, P_*dat-1*_::hVPS41^S285P^ + P_*dat-1*_::hVPS41^R662*^ (baEx217] in the UA44 (UA391) or BY250 (UA388) backgrounds was analyzed. For all strains, three representative lines of each were selected for further analysis.

### Dopaminergic neurodegeneration analysis in *C. elegans*

*C. elegans* dopaminergic neurons were analyzed for degeneration as previously described(32,33). Briefly, synchronized progeny from the BY250, UA44, and experimental hVPS41 variant strains were produced from a 3-hour egg-lay and grown at 20°C. Animals were analyzed at days 7 and 10 post-hatching (4- and 7-day-old adults). On the day of analysis, the 6 anterior dopaminergic neurons [4 CEP (cephalic) and 2 ADE (anterior deirid)] were examined in 30 adult hermaphrodite worms, which were immobilized on glass cover slips using 3mM levamisole and transferred onto 2% agarose pads on microscope slides.

In quantifying neurodegeneration, animals are scored as “normal” when a full complement of all 6 anterior dopaminergic neurons are present and the neuronal processes are fully intact, as previously reported(33,85,86). In total, at least 90 adult worms (30 worms/trial x 3 trials = 90 total animals) were analyzed for each independent strain and at least 270 adult worms were analyzed for each independent transgenic strain (30 worms/trial x 3 independent transgenic lines x 3 trials = 270 total animals/construct). Values of each independent experiment were normalized to the BY250 control and representative analyses were averaged and statistically analyzed by Prism and One-Way ANOVA.

### Protein purification

The appendage domain of rat AP-3 delta were amplified by PCR and cloned in-frame into pET15b-MBP-MCS. The recombinant proteins were induced overnight at 20°C in Escherichia coli BL21 (DE3) using 0.1 mM IPTG. The bacteria were lysed by sonication in in 50 mM Tris, 300mM NaCl, pH 8.0 and MBP fusions purified by incubation with amylose resin beads (NEB) and eluted with 50mM maltose in 50 mM Tris, 300mM NaCl, pH 8.0. Two days after infection with baculoviruses produced according to the manufacturer’s instructions (Invitrogen), Hi5 cells were harvested by centrifugation and resuspended in binding buffer (50 mM Tris, 300 mM NaCl, pH 8.0) with 1 mM PMSF and protease inhibitors (Complete, Roche). The cells were lysed by sonication and pelleted for 20 min at 21,000g using a 4°C Eppendorf 5424 centrifuge. Supernatant was passed through a 0.2 μm filter and incubated with Co-Nta agarose (GoldBio) for 1 hr at 4°C. The agarose was then washed five times in binding buffer with 10-40 mM imidazole and His-hVPS41 eluted with 500 mM imidazole. The eluates were immediately desalted on a Secadex-5 column (Prometheus) into lysis buffer. 50ug of MBP or MBP-AP3 were loaded onto amylose-resin for 30min at 4C. Unbound protein was removed by several washing steps, and varying amount of VPS41 were incubated with the MBP resin to assess interaction. Bound protein was eluted and evaluated by quantitative fluorescence immunoblotting.

### Secretion Assays

PC12 cells were plated on poly-L-lysine, washed, and incubated in Tyrode’s buffer containing 2.5 mM K+ (basal) or 90 mM K+ (stimulated) for 30 min at 37°C. The supernatant was collected, cell lysates prepared as previously described(30), and samples were analyzed by quantitative fluorescence immunoblotting.

## Statistics

Quantification of all western blots was performed using Fiji software(80). Two-tailed Students *t* tests were performed to compare two samples whereas ANOVA was performed for comparison of multiple samples to analyze statistical significance. Data distribution was assumed to be normal, but this was not formally tested. All error bars represent the Standard Error of the Mean (SEM).

## Author contributions

RW, CB, CB, LC, FZ, SZ, EG and TV carried out the experiments. RJ, AF, SB, CS, AL, GY and DC performed clinical investigation on both patients. RW, CB, PS, JB, NL, GC, CA, KC, DC and JK contributed to the study design. AF, FZ, JB, NL, SB, CA, GC, KC and DC reviewed and revised the manuscript. CA, GC, KC, DC and JK designed and supervised the study. JK acquired funding. RW and JK wrote the manuscript.

## Acknowledgements

We thank our colleagues of the Center for Molecular Medicine for fruitful discussions. We would like to thank Bas van Zuijlen for the generation of the VPS11 and VPS18 knockout cell lines. We thank Dr. Stephanie de Poot for kindly providing the APEX2 constructs, and Prof. R. Roepman and Prof. J.J.C. Neefjes for providing the GFP-pcDNA-Dest53 and pEGFP-C1-mCherry-VPS41 constructs respectively. Catherine Rabouille (Hubrecht, The Netherlands) is acknowledged for her critical input throughout our studies. We thank Prof. Dr. C.M.A Ravenswaaij-Aarts and I. Stolte-Dijkstra for proving information and patient fibroblasts from the third VPS41 patient. Support for the *C. elegans* studies came from a grant from the National Institutes of Health [R15 NS075684-01 to G.A.C.]. N.L. is supported by a ZonMW TOP grant [40-00812-98-16006 to J.K]. P.S is supported by a DFG grant [FOR2625 to J.K. as part of the Research Consortium].

## References

1. Saftig P, Klumperman J. Lysosome biogenesis and lysosomal membrane proteins: trafficking meets function. Nat Rev Mol Cell Biol. 2009;10(9):623–35.

2. Luzio JP, Pryor PR, Bright NA. Lysosomes: fusion and function. Nat Rev. 2007;8:622–32.

3. Settembre C, Fraldi A, Medina DL, Ballabio A, Children T. Signals for the lysosome: a control center for cellular clearance and energy metabolism. Nat Rev Mol Cell Biol. 2015;14(5):283–96.

4. Richardson DR, et al. Mitochondrial iron trafficking and the integration of iron metabolism between the mitochondrion and cytosol. Proc Natl Acad Sci. 2010;107(24):10775–82.

5. Mony VK, Benjamin S, O’Rourke EJ. A lysosome-centered view of nutrient homeostasis. Autophagy. 2016;12(4):619–31.

6. Pols MS, ten Brink C, Gosavi P, Oorschot V, Klumperman J. The HOPS proteins hVps41 and hVps39 are required for homotypic and heterotypic late endosome fusion. Traffic. 2013;14(2):219–32.

7. Caplan S, Hartnell LM, Aguilar RC, Naslavsky N, Bonifacino JS. Human Vam6p promotes lysosome clustering and fusion in vivo. J Cell Biol. 2001;7:109–21.

8. Van der Kant R, et al. Characterization of the mammalian CORVET and HOPS complexes and their modular restructuring for endosome specificity. J Biol Chem. 2015;290(51):30280–90.

9. Wurmser AE, Sato TK, Emr SD, Vam V. New Component of the Vacuolar Class C-Vps Complex Couples Nucleotide Exchange on the Ypt7 GTPase to SNARE-dependent Docking and Fusion. J Cell Biol. 2000;151(3):551–62.

10. Seals DF, Eitzen G, Margolis N, Wickner WT, Price A. A Ypt/Rab effector complex containing the Sec1 homolog Vps33p is required for homotypic vacuole fusion. Proc Natl Acad Sci. 2000;97(17):9402–7.

11. Beek J Van Der, Jonker C, Welle R Van Der, Liv N, Klumperman J. CORVET, CHEVI and HOPS – multisubunit tethers of the endo-lysosomal system in health and disease. J Cell Sci. 2019;132

12. Radisky DC, Snyder WB, Emr SD, Kaplan J. Characterization of VPS41, a gene required for vacuolar trafficking and high-affinity iron transport in yeast. Cell Biol. 1997;94:5662–6.

13. Nickerson DP, Brett CL, Merz AJ. Vps-C complexes: gatekeepers of endolysosomal traffic. Curr Opin Cell Biol. 2009;21(4):543–51.

14. Aoyama M, Sun-Wada GH, Yamamoto A, Yamamoto M, Hamada H, Wada Y. Spatial Restriction of Bone Morphogenetic Protein Signaling in Mouse Gastrula through the mVam2-Dependent Endocytic Pathway. Dev Cell. 2012;22(6):1163–75.

15. Lin X, et al. RILP interacts with HOPS complex via VPS41 subunit to regulate endocytic trafficking. Sci Rep. 2014;4:7282.

16. Khatter D, Raina VB, Dwivedi D, Sindhwani A, Bahl S, Sharma M. The small GTPase Arl8b regulates assembly of the mammalian HOPS complex on lysosomes. J Cell Sci. 2015;128(9):1746–61.

17. Takáts S, et al. Interaction of the HOPS complex with Syntaxin 17 mediates autophagosome clearance in Drosophila. 2014;25(8):1338–54.

18. Pols MS, et al. hVps41 and VAMP7 function in direct TGN to late endosome transport of lysosomal membrane proteins. Nat Commun. 2013;4:1361.

19. Swetha MG, et al. Lysosomal membrane protein composition, acidic pH and sterol content are regulated via a light-dependent pathway in metazoan cells. Traffic. 2011;12(8):1037–55.

20. Cowles CR, Odorizzi G, Payne GS, Emr SD. The AP-3 Adaptor Complex Is Essential for Cargo-Selective Transport to the Yeast Vacuole. Cell. 1997;91:109–18.

21. Merz CGA and AJ. HOPS Interacts with Apl5 at the Vacuole Membrane and Is Required for Consumption of AP-3 Transport Vesicles. Mol Biol Cell. 2009;20(22):4042–56.

22. Cabrera M, et al. Phosphorylation of a membrane curvature-sensing motif switches function of the HOPS subunit Vps41 in membrane tethering. J Cell Biol. 2010;191(4):845–59.

23. Darsow T, Katzmann DJ, Cowles CR, Emr SD. Vps41p function in the alkaline phosphatase pathway requires homo-oligomerization and interaction with AP-3 through two distinct domains. Mol Biol Cell. 2001;12(1):37–51.

24. Rehling P, et al. Formation of AP-3 transport intermediates requires Vps41 function. Nat Cell Biol. 1999;1(6):346–53.

25. Hummer BH, et al. HID-1 controls formation of large dense core vesicles by influencing cargo sorting and trans-Golgi network acidification. Mol Biol Cell. 2017;28(26):3870–80.

26. Orci L, et al. The trans-most cisternae of the Golgi complex: A compartment for sorting of secretory and plasma membrane proteins. Cell. 1987;51(6):1039–51.

27. Tooze SA, Huttner WB. Cell-free sorting to the regulated and constitutive pathways. Cell. 1990;60:837–47.

28. Eaton B a, Haugwitz M, Lau D, Moore HP. Biogenesis of regulated exocytotic carriers in neuroendocrine cells. J Neurosci. 2000;20(19):7334–44.

29. Burgess TL and Kelly RB. Constitutive and Regulated Secretion of proteins. Ann Rev Cell Biol. 1987;3:243–93.

30. Asensio CS, Sirkis DW, Edwards RH. RNAi screen identifies a role for adaptor protein AP-3 in sorting to the regulated secretory pathway. J Cell Biol. 2010;191(6):1173–87.

31. Asensio CS, et al. Self-Assembly of VPS41 Promotes Sorting Required for Biogenesis of the Regulated Secretory Pathway. Dev Cell. 2013;27(4):425–37.

32. Harrington AJ, Yacoubian TA, Slone SR, Caldwell KA, Caldwell GA. Functional analysis of VPS41-mediated neuroprotection in Caenorhabditis elegans and mammalian models of Parkinson’s disease. J Neurosci]. 2012;32(6):2142–53.

33. Hamamichi S, Rivas RN, Knight AL, Cao S, Caldwell KA, Caldwell GA. Hypothesis-based RNAi screening identifies neuroprotective genes in a Parkinson’s disease model. PNAS. 2008;105(2):728–33.

34. Ruan Q, Harrington AJ, Caldwell KA, Caldwell GA, Standaert DG. VPS41, a protein involved in lysosomal trafficking, is protective in Caenorhabditis elegans and mammalian cellular models of Parkinson’s disease. Neurobiol Dis. 2010;37(2):330–8.

35. Griffin EF, Yan X, Caldwell KA, Caldwell GA. Distinct functional roles of Vps41-mediated neuroprotection in Alzheimer’s and Parkinson’s disease models of neurodegeneration. Hum Mol Genet. 2018;27(24):4176–93.

36. Ibarrola-Villava M, et al. Genes involved in the WNT and vesicular trafficking pathways are associated with melanoma predisposition. Int J Cancer. 2015;136(9):2109–19.

37. Papa V, et al. The role of ultrastructural examination in storage diseases. Ultrastruct Pathol. 2010;34(5):243–51.

38. Peden AA, Oorschot V, Hesser BA, Austin CD, Scheller RH, Klumperman J. Localization of the AP-3 adaptor complex defines a novel endosomal exit site for lysosomal membrane proteins. J Cell Biol. 2004;1065–76.

39. Pols MS, et al. hVps41 and VAMP7 function in direct TGN to late endosome transport of lysosomal membrane proteins. Nat Commun. 2013;4:1312–61.

40. Klumperman J, Raposo G. The complex ultrastructure of the endolysosomal system. Cold Spring Harb Perspect Biol. 2014;6(10).

41. Huotari J, Helenius A. Endosome maturation. EMBO J. 2011;30(17):3481–500.

42. Hunter M, Scourfield EJ, Emmott E, Graham SC. VPS18 recruits VPS41 to the human HOPS complex via a RING-RING interaction. Biochem J. 2017; 474(21):3615–3626.

43. Jia R, Guardia CM, Pu J, Chen Y, Bonifacino JS. BORC coordinates encounter and fusion of lysosomes with autophagosomes. Autophagy. 2017;13(10):1648–63.

44. Adzhubei IA, et al. A method and server for predicting damaging missense mutations. Nat Methods. 2010;7(4):248–9.

45. Jiang P, et al. The HOPS complex mediates autophagosome-lysosome fusion through interaction with syntaxin 17. Mol Biol Cell. 2014;25(8):1327–37.

46. Nakamura S, Yoshimori T. New insights into autophagosome – lysosome fusion. J Cell Sci. 2017;1209–16.

47. McEwan DG, et al. PLEKHM1 regulates autophagosome-lysosome fusion through HOPS complex and LC3/GABARAP proteins. Mol Cell. 2015;57(1):39–54.

48. Kimura S, Noda T, Yoshimori T. Dissection of the Autophagosome Maturation Process by a Novel Reporter Protein, Tandem Fluorescent-Tagged LC3. Autophagy. 2007;3(5):452–60

49. Palmieri M, et al. Characterization of the CLEAR network reveals an integrated control of cellular clearance pathways. Hum Mol Genet. 2011;20(19):3852–66.

50. Martina JA, et al. The nutrient-responsive transcription factor TFE3 promotes autophagy, lysosomal biogenesis, and clearance of cellular debris. Sci Signal. 2014;7(309):1–16.

51. Settembre C, et al. TFEB Links Autophagy to Lysosomal Biogenesis. Science. 2011;332(6036):1429–33.

52. Cabrera M, et al. Phosphorylation of a membrane curvature–sensing motif switches function of the HOPS subunit Vps41 in membrane tethering. J Cell Biol. 2010;191(4):845–59.

53. Reich SG and Savitt JM. Parkinson’s Disease. Med Clin North Am. 2019;103(2):337–350.

54. Jankovic J. Parkinson’s disease: clinical features and diagnosis. J Neurol Neurosurg Psychiatry. 2008;79(4):368–76.

55. Laplante M, Sabatini DM. mTOR Signaling in Growth Control and Disease. Cell. 2012;149(2):274–93.

56. Kim YC, Guan K. mTOR: a pharmacologic target for autophagy regulation. 2015;125(1):25–32.

57. Raben N, Puertollano R. TFEB and TFE3: Linking Lysosomes to Cellular Adaptation to Stress. Annu Rev Cell Dev Biol. 2016;32:255–278.

58. Mauthe M, et al. Chloroquine inhibits autophagic flux by decreasing autophagosome-lysosome fusion. Autophagy. 2018;14(8):1435–55.

59. Ebrahimi-Fakhari D, Wahlster L, Hoffmann GF, Kölker S. Emerging role of autophagy in pediatric neurodegenerative and neurometabolic diseases. Pediatr Res. 2014;75(1–2):217–26.

60. Flinn RJ, Yan Y, Goswami S, Parker PJ, Backer JM. The Late Endosome is Essential for mTORC1 Signaling. Mol Biol Cell. 2010;21:833–41.

61. Slade L, Pulinilkunnil T. The MiTF/TFE Family of Transcription Factors: Master Regulators of Organelle Signaling, Metabolism and Stress Adaptation. Mol Cancer Res. 2017;15(12):1637–1643.

62. Sardiello M, et al. A Gene Network Regulating Lysosomal Biogenesis and Function. Science. 2009;325(5939):473–7.

63. Stadel D, et al. TECPR2 Cooperates with LC3C to Regulate COPII-Dependent ER Export. Mol Cell. 2015;60(1):89–104.

64. Cowles CR, Snyder WB, Burd CG, Emr SD. Novel Golgi to vacuole delivery pathway in yeast: Identification of a sorting determinant and required transport component. EMBO J. 1997;16(10):2769–82.

65. Mazzulli JR, et al. Gaucher Disease Glucocerebrosidase and a-synuclein form a bidirectional pathogenic loop in synucleinopathies. Cell. 2011;146(1):37–52.

66. Do J, Mckinney C, Sharma P, Sidransky E. Glucocerebrosidase and its relevance to Parkinson disease. Mol Neurodegener. 2019;14(1):36.

67. Zhen Y, Li W. Impairment of autophagosome-lysosome fusion in the buff mutant mice with the VPS33A(D251E) mutation. Autophagy. 2015;11(9):1608–22.

68. Edvardson S, et al. Hypomyelination and developmental delay associated with VPS11 mutation in Ashkenazi-Jewish patients. J Med Genet. 2015;52(11):749–53.

69. Hörtnagel K, et al. The second report of a new hypomyelinating disease due to a defect in the VPS11 gene discloses a massive lysosomal involvement. J Inherit Metab Dis. 2016;39(6):849–57.

70. Zhang J, et al. A Founder Mutation in VPS11 causes an autosomal recessive leukoencephalopathy linked to autophagic defects. PLOS Genet. 2016;12(4):e1005848.

71. Kondo H, et al. Mutation in VPS33A affects metabolism of glycosaminoglycans: a new type of mucopolysaccharidosis with severe systemic symptoms. Hum Mol Genet. 2016;26(1):173–183.

72. Chintala S, et al. The VPS33a gene regulates behavior and cerebellar Purkinje cell number. Brain Res. 2009;1266:18–28.

73. Dursun A, et al. A probable new syndrome with the storage disease phenotype caused by the VPS33A gene mutation. Clin Dysmorphol. 2017;26(1):1–12.

74. Pavlova EV, et al. The lysosomal disease caused by mutant VPS33A. Hum Mol Genet. 2019;28(15):2514–30.

75. Lam HC, et al. p62/SQSTM1 cooperates with hyperactive mTORC1 to regulate glutathione production, maintain mitochondrial integrity and promote tumorigenesis. Cancer res. 2017;77(12):3255–67.

76. An W, Cowburn RF, Li L, Braak H, Alafuzoff I, Iqbal K. Up-regulation of phosphorylated / activated p70 S6 Kinase and its relationship to neurofibrillary pathology in Alzheimer’s Disease. Am J Pathol. 2003;163(2):591–607.

77. Wong M. Mammalian Target of Rapamycin (mTOR) pathways in neurological diseases. Biochem J. 2013;85(0 1):1–27.

78. Edvardson S, et al. Hypomyelination and developmental delay associated with VPS11 mutation in Ashkenazi-Jewish patients. J Med Genet. 2015 Nov;52(11):749–53.

79. Cai X, et al. Homozygous mutation of VPS16 gene is responsible for an autosomal recessive adolescent-onset primary dystonia. Sci Rep. 2016;6(1):25834.

80. Schindelin J, et al. Fiji: an open-source platform for biological-image analysis. Nat Methods. 2012;9(7):676–82.

81. Slot JW, Geuze HJ. Cryosectioning and immunolabeling. Nat Protoc. 2007;2(10):2480–91.

82. Liou W, Geuze HJ, Slot JW. Improving structural integrity of cryosections for immunogold labeling. Histochem Cell Biol. 1996;106(1)41–58.

83. Ran FA, Hsu PD, Wright J, Agarwala V, Scott DA, Zhang F. Genome engineering using the CRISPR-Cas9 system. Nat Protoc. 2013;8(11):2281–308.

84. Haeussler M, et al. Evaluation of off-target and on-target scoring algorithms and integration into the guide RNA selection tool CRISPOR. Genome Biol. 2016;17(1):148.

85. Cao S, Gelwix CC, Caldwell KA, Caldwell GA. Torsin-mediated protection from cellular stress in the dopaminergic neurons of Caenorhabditis elegans. Neurobiol Dis. 2005;25(15):3801–12.

86. Harrington AJ, Hamamichi S, Caldwell GA, Caldwell KA. C. elegans as a model organism to investigate molecular pathways Involved with Parkinson’s Disease. Dev Dyn. 2010;239(5):1282–95.

