## Supplemental information and figures for "*VPS41* recessive mutation causes ataxia and dystonia with retinal dystrophy and mental retardation by inhibiting HOPS function and mTORC1 signaling"

**Clinical presentation**

**Patient 1**

Currently 20 years-old male was born to healthy and non-consanguineous parents. His birth weight, length and head circumference (OFC) were within the normal range. At approximately two months of age, he was noted to have poor fixing and following and ophthalmological examination showed hypopigmentation of the retina and narrow optic nerve suggestive of retinal dystrophy. Electroretinogram (ERG) tracings were markedly abnormal but gradually improved and are currently normal. As an infant he had hypotonia and global developmental delay. His hand-coordination was poor and he was diagnosed with autism spectrum disorder. Extensive metabolic studies showed no detectable abnormalities and brain MRI initially showed only mild hypomyelination. However, brain MRIs showed progressive hypoplasia of the corpus callosum (CC) and cerebellum (Figure 1A). By approximately six months he was noted to have low muscle tone, delayed fine and gross motor skills and marked tremor which further impaired his fine motor skills. His deep tendon reflexes (DTRs) were markedly impaired to absent and his plantars were extensor. There was upper extremity tremor, pass-pointing, significant ataxia of the lower limbs and spastic ataxia on supported gait.

At age 7 he had severe global developmental delay with remarkable tremor. His language skills were impaired but less than his gross and fine motor skills.

On examination at age 20 he had global developmental delay, generalized hypotonia, bilateral flat feet with a mild feet dorsiflexion and absence of all DTRs with bilateral flexor plantar response. He had bilateral limb dysmetria and ataxic gait. He had neither tremor nor dystonia during gait but there was an abnormal posturing in the upper and lower extremities. He had repetitive behaviors, pervasive tendencies and some obsessive behaviors (video 1). His karyotype was normal and male (46, XY) and microarray analysis showed a de novo duplication of 2.568Mb at 1q21.1 of unknown significance and a deletion of 0.293Mb at 12q14.3 of maternal origin. Whole-exome sequencing revealed compound heterozygote variants in the VPS41 gene.

**Patient 2**

The younger brother, the male sibling of patient 1 was born two years later. His birth weight, length and OFC were within the normal range. Ophthalmological examination including ERG at 3 months of age noted mild retinal dystrophy with possible hyperopia which improved over the years. Hypotonia was noted in infancy but with no latching difficulties. His fine motor skills were delayed due to bilateral tremor which significantly worsened with excitement. At 3 years he was noted to have global developmental delay and he had hyperacusis. Extraocular movements were full but visual pursuit was saccadic. DTRs were absent in the upper and lower limbs and the plantars were extensor bilaterally.

By 5 years of age he was noted to have dystonic postures of both lower limbs as well as ataxic gait. The dystonia spread to his upper limbs, trunk and neck within the following year and further worsened compromising his ability to ambulate by age 7. A year later, tongue movement and feeding dystonia were also noted. Initial brain MRI showed mild hypomyelination. However, brain MRI at 5 years showed mild hypoplasia of the cerebellum and cerebellar vermis and a thin corpus callosum. The brain MRI was repeated at the age of 10 and confirmed the thin corpus callosum and mild progression of the cerebellar atrophy. This MRI also showed bilateral hypointensity in the globus pallidus (GP) compatible with early iron deposition, as well as abnormal hyperintensity in the dentate nucleus and cerebellar cortex. Nerve conduction studies were normal and CSF analysis showed normal protein, glucose, cytology and neurotransmitters (pyridoxal phosphate, 5-methyltetrahydrofolate, tetrahydropterin, neopterin, succinyl adenosine). Analyses of genes known to cause X-linked mental retardation and dystonia as well as genes associated with iron deposition in the basal ganglia were tested but no mutations were found (Table S1). Microarray analysis showed a single copy loss of a deletion of 0.293Mb at 12q14.3 of unknown significance, of maternal origin. Whole exome sequencing revealed compound heterozygote variants in the VPS41 gene.

Levodopa, trihexyphenidyl, clonazepam, baclofen as well as botulinum toxin injections at the highest tolerated doses were all ineffective in controlling his generalized dystonia. At 12.5 years he was referred for deep brain stimulation (DBS) of the GP pars interna (GPi). Examination at that time revealed generalized dystonia involving his face, tongue, neck, trunk and four limbs. There was neither spasticity nor pyramidal signs and parkinsonism, tremor or myoclonus identified. Cerebellar signs and deep tendon reflexes were hard to assess due to his severe dystonia (Video 2). Bilateral GPi DBS was performed at the age of 13 improving trunk, head and arm thus allowing him to sit straight and gain fine motor skills with both hands. By contrast, lower limbs dystonia responded only partially to DBS, requiring complementary treatment with botulinum toxin with slight and fluctuating benefit. The improvements with DBS have been stable till the last follow up at 18 years [4.5 years after DBS (Video 3)].

### Supplemental table 1

**Table 1.** Mutation analysis on several genes associated with X-linked mental retardation and dystiona as well as iron deposition in the basla ganglia were performed. No mutations were found.

| | **Tested gene** |
|---|---|
| DYT1 | TOR1A |
| Dystonia Myoclonus, DYT11 | SGCE |
| FAHN, spastic paraplegia 35 | FA2H |
| Infantile neuroaxonal dystrophyl, Neurodegeneration with brain iron accumulation 2B | PLA2G6 |
| Partington syndrome | ARX |
| Renpenning syndrome | PQBP1 |
| syndromic X-linked mental retardation, Claes-Jensen type | JARID1C |
| Mental retardation, X-linked 58 | TM4SF2 |
| Mental retardation, X-linked 63 | FACL4 |
| Mental retardation, X-linked 89 | ZNF41 |
| Mental retardation, X-linked 90 | DLG3 |
| Rett Syndrome | MECP2 |

### Supplementary data

#### Figure S1

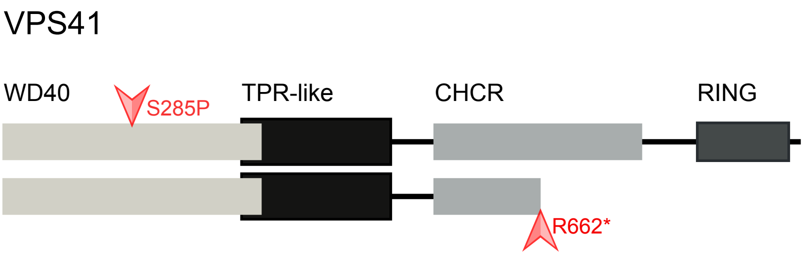


#### Figure S2

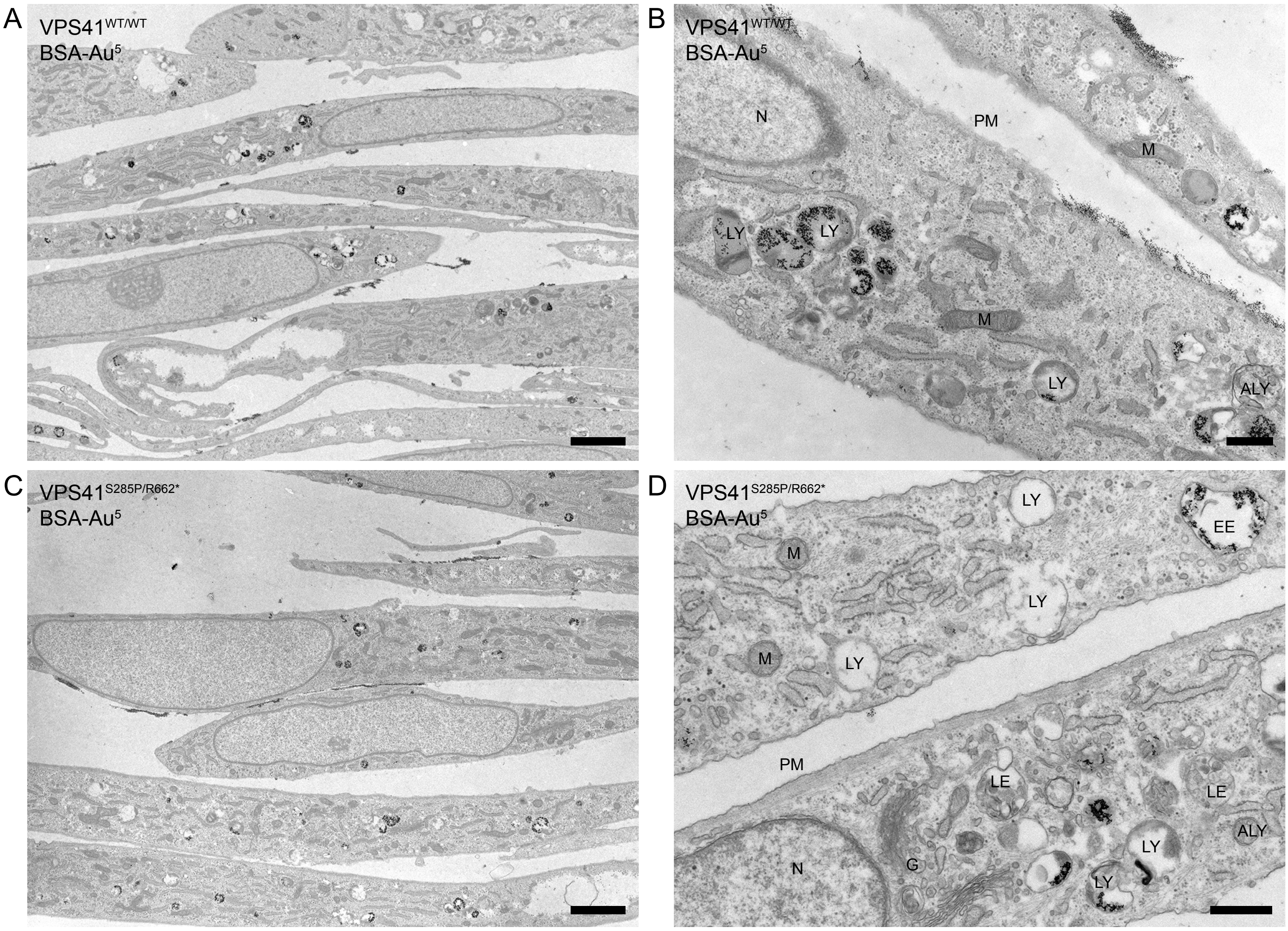

**Figure S1**. **Mutations in VPS41.** Outline of the VPS41 protein depicting distinct domains. The WD40 domain facilitates protein-protein interactions, the CHCR and RING domains enable homo-oligomerization and are required for HOPS complex formation and regulated secretion. Mutations in *VPS41* were identified using exome sequencing. Both patients bear compound heterozygous mutations; a missense mutation $VPS41^{S285P}$ in the WD40 domain and a nonsense mutation $VPS41^{R662*}$ at the C-terminus resulting in a premature stopcodon.

**Figure S2. Mutations in *VPS41* do not cause a lysosomal storage phenotype.** Electron micrographs of $VPS41^{WT/WT*}$(**A,B**), and $VPS41^{285P/R662*}$ (**C,D**), fibroblasts. Cells were incubated with BSA-Au[5] for 2 hours to visualize endo-lysosomal compartments. Both primary fibroblast cell lines show a high variation in the appearance of endo-lysosomal compartments. There is no aberrant swelling of endo-lysosomal organelles in patient $VPS41^{S285P/R662*}$ fibroblasts. ALY= Autolysosome, G= Golgi, LE= Late endosome, LY= Lysosome, M= Mitochondria, N= Nucleus, PM= Plasma membrane. Scale bars A, C: 2µm, scale bars B, D: 500nm.

Figure S3

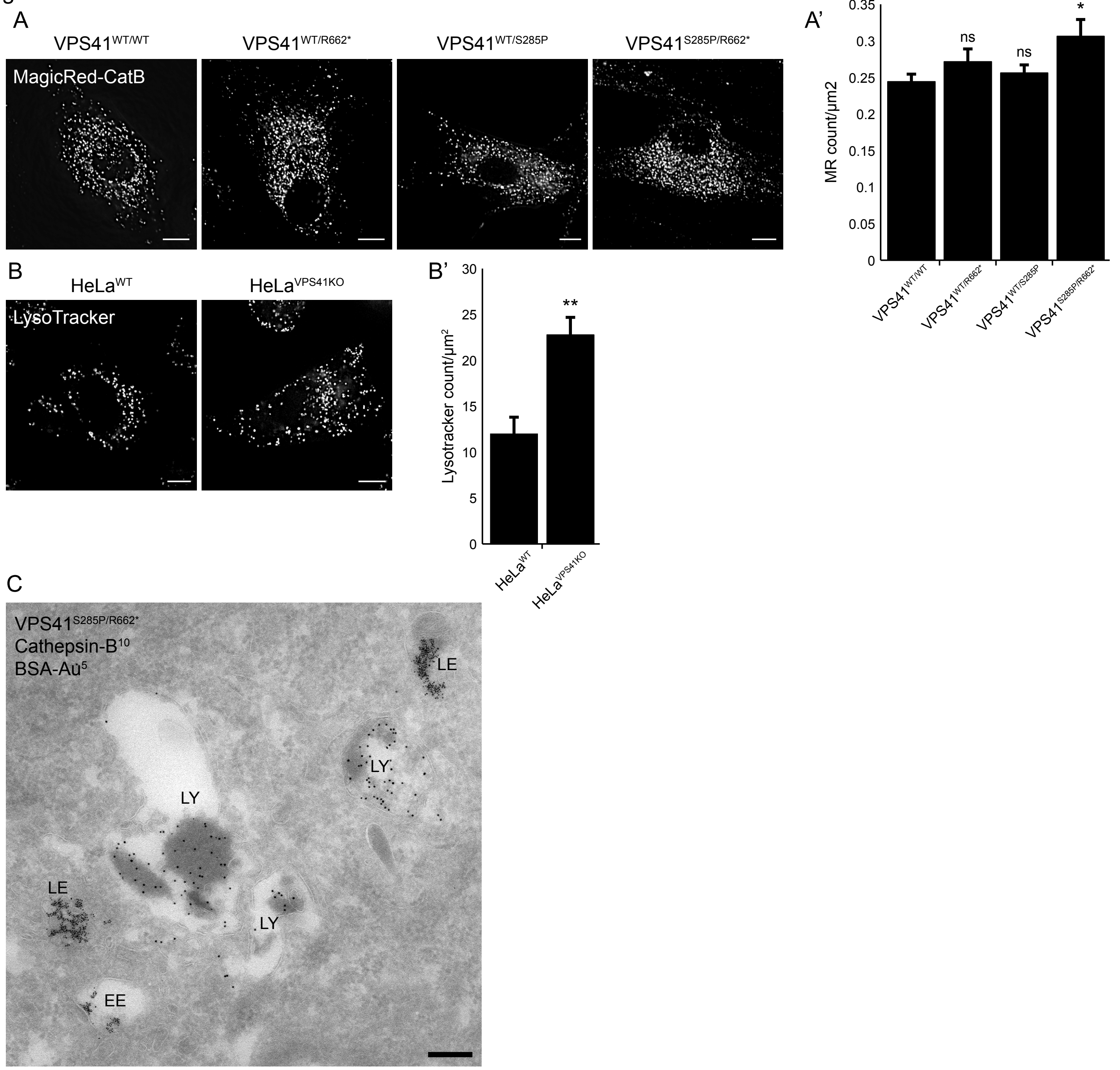


Figure S4

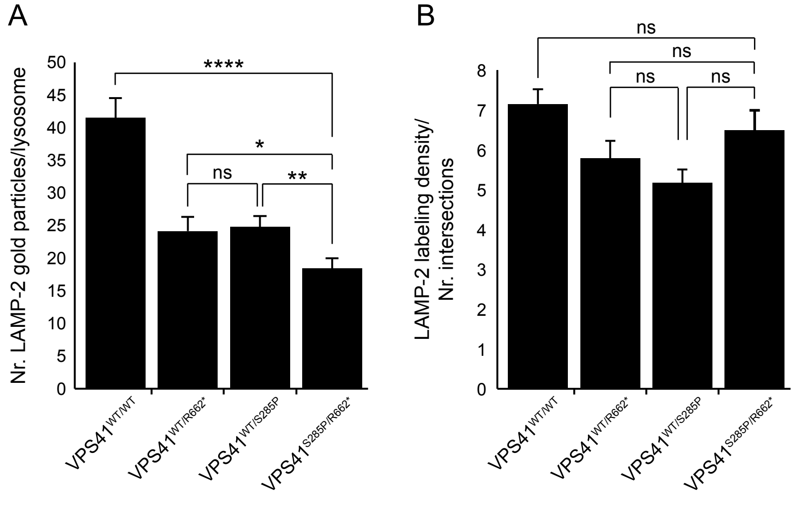

**Figure S3. Mutant VPS41 or VPS41$^{KO}$ results in increased numbers of enzymatically active, acidified compartments. (A)** *VPS41$^{WT/WT}$*, *VPS41$^{WT/S285P}$*, *VPS41$^{WT/R662*}$* and *VPS41S$^{285P/R662*}$* fibroblasts incubated with MagicRed-cathepsin B and imaged by fluorescence microscopy to visualize enzymatically active compartments. *VPS41$^{S285P/R662*}$* fibroblasts show significantly more cathepsin B active compartments (quantified in **S3A'**). (n=3). Bars 10µm. Error bars represent the SEM. *p<0.05; one-ANOVA with Bonferroni correction for multiple comparisons. **(B)** HeLa$^{WT}$ and HeLa$^{VPS41KO}$ cells incubated with LysoTrackerRed to assess the prevalence of acidified compartments in the absence of VPS41. Hela$^{VPS41KO}$ cells show a 2-fold increase in number of acidified compartments (quantified in **S3B'**). (n=3) Bars 10µm. Error bars represent the SEM. Unpaired *t* test, **p<0.01. **(C)** Immuno-electron microscopy of *VPS41$^{S285P/R662*}$* fibroblasts loaded for 2h with BSA conjugated to 5 nm gold (BSA-Au$^5$) and labeled for cathepsin B (10 nm gold particles). Lysosomes are positive for cathepsin B, indicating that patient fibroblasts have fully equipped lysosomes. EE= Early endosome, LE= late endosome, LY= lysosome. Scale bar: 200nm.

**Figure S4. Lysosomes of patient fibroblasts are reached by LAMP-2. (A)** To determine if lysosomes in patient derived fibroblasts are reached by LAMP-2, *VPS41$^{WT/WT}$*, *VPS41$^{WT/S285P}$*, *VPS41$^{WT/R662*}$* and *VPS41$^{S285P/R662*}$* fibroblasts were prepared for immuno-EM and labeled for LAMP-2. The number of LAMP-2 representing gold particles was counted in each LAMP-2 positive lysosome. *VPS41$^{S285PR662*}$* have significantly less LAMP-2. Error bars represent the SEM. *p<0.05, **p<0.01, ****p<10$^{-5}$; one-ANOVA with Tukey correction for multiple comparisons. **(B)** After correction for lysosome size, measured by number of intersections of a grid (200nm grid size) covering a specific lysosome, the density of LAMP-2 label showed no significant difference between *VPS41$^{WT/WT}$*, *VPS41$^{WT/S285P}$*, *VPS41$^{WT/R662*}$* or *VPS41$^{S285P/R662*}$* fibroblasts. Error bars represent the SEM; one-ANOVA with Tukey correction for multiple comparisons.

Figure S5

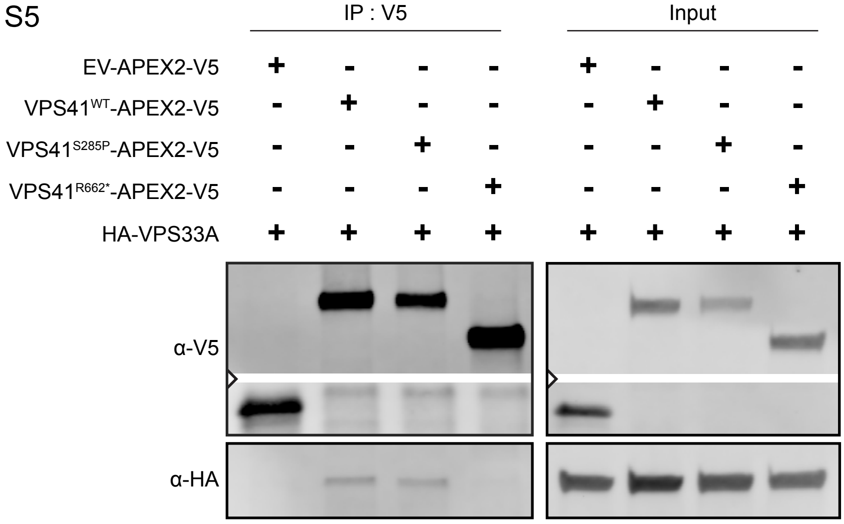


Figure S6

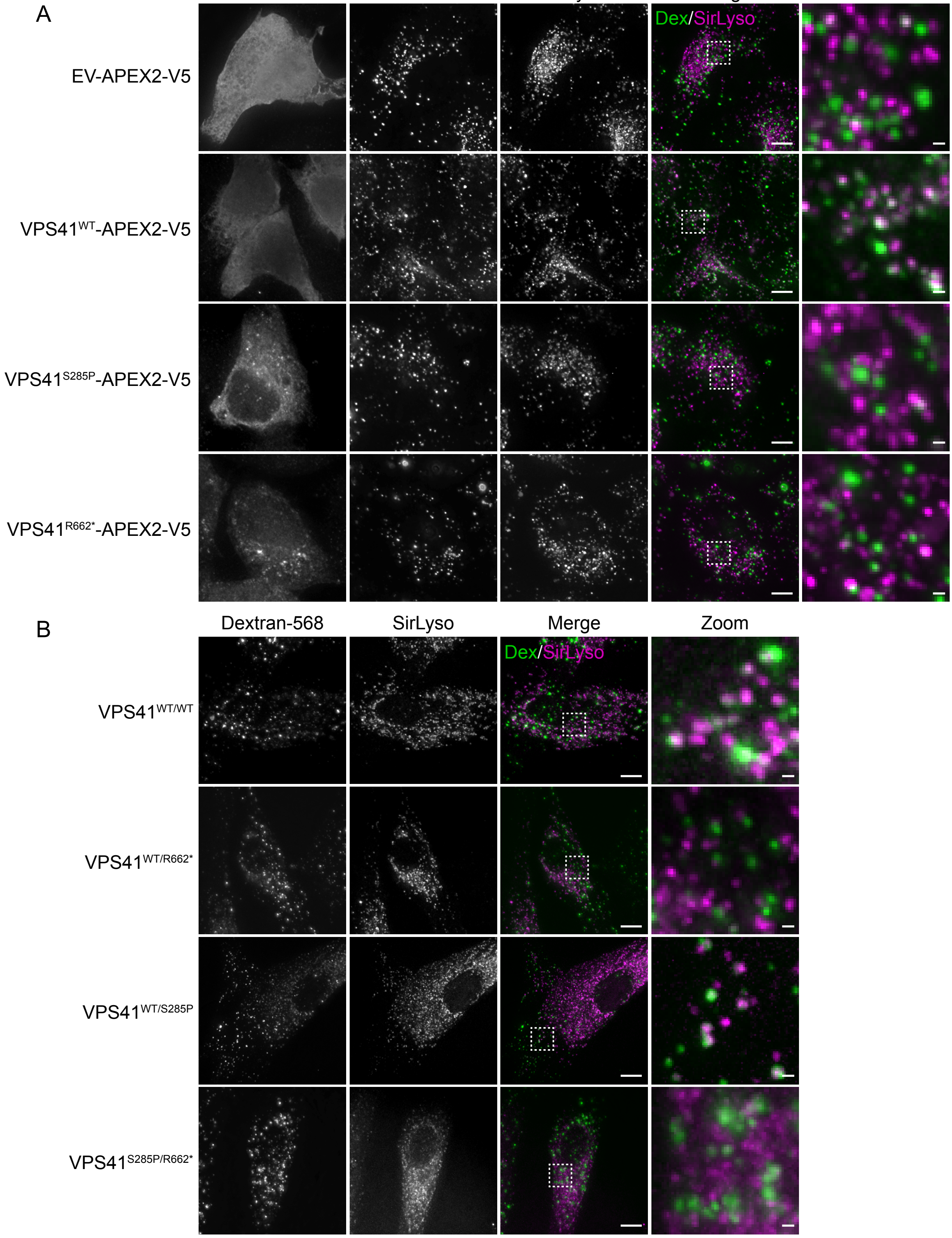

**Figure S5. VPS41$^{R662*}$ does not interact with HOPS subunit VPS33A. (A)** Immunopreciptation (IP) on HeLa$^{WT}$ cells co-expressing VPS41$^{WT}$-APEX2-V5, VPS41$^{S285P}$-APEX2-V5 or VPS41$^{R662*}$-APEX2-V5 and HA-VPS33A. Western blot analysis shows that in contrast to VPS41$^{WT}$ and VPS41$^{s285P}$ the VPS41$^{R662*}$ variant does not interact with VPS33A. n=3.

**Figure S6. Loss of VPS41$^{WT}$ results in impaired HOPS dependent late endosome – lysosome fusion. (A)** HeLa$^{VPS41KO}$ cells transfected with VPS41$^{WT}$-APEX2-V5 showed a significant increase in colocalization between Dextran and SiR-Lysosome cathepsin D, indicating rescue of the endocytosis phenotype observed in *VPS41*$^{KO}$ cells. Expression of either of the *VPS41* variant resulted in poor colocalization, indicating that both variants fail to rescue HOPS complex functionality. Bars 10µm, zoom 1µm. **(B)** *VPS41*$^{WT/WT}$, VPS41$^{WT/S285P}$. *VPS41*$^{WT/R662*}$ and *VPS41*$^{S285P/R662*}$ primary fibroblasts were incubated with Dextran-ALEXA568 and SiR-Lysosome cathepsin D for 2h and 3h respectively. Co-localization representing delivery of Dextran to enzymatically active lysosomes was decreased in *VPS41*$^{WT/R662*}$ and *VPS41*$^{S285P/R662*}$ cells, indicating reduced fusion efficiency between late endosomes and lysosomes. Bars 10µm, zoom 1µm.

Figure S7

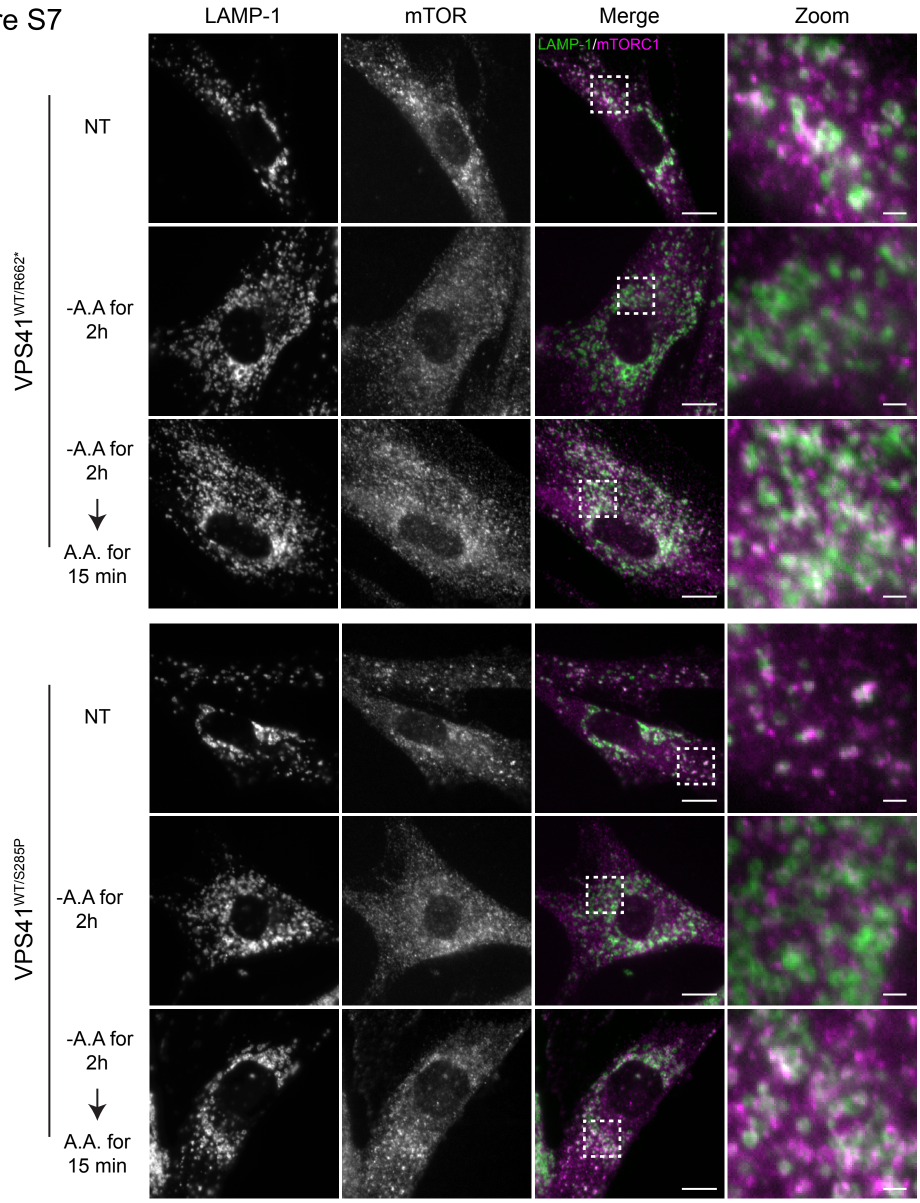

LAMP-1
mTOR
Merge
Zoom
LAMP-1/mTORC1
VPS41WT/R662*
NT
-A.A for 2h
-A.A for 2h
A.A. for 15 min
VPS41WT/S285P
NT
-A.A for 2h
-A.A for 2h
A.A. for 15 min

**Figure S7. Paternal (VPS41$^{WT/S285P}$) and maternal (VPS41$^{WT/R662*}$) show proper mTORC1 localization in response to nutrient availability.** Immunofluorescence of parental fibroblasts labeled for LAMP-1 and mTORC1. *VPS41*$^{WT/S285P}$ and *VPS41*$^{WT/R662*}$ show an appropriate mTORC1 response upon nutrient deprivation (-Amino Acids (-AA)) or stimulation (-AA, +AA).

Figure S8

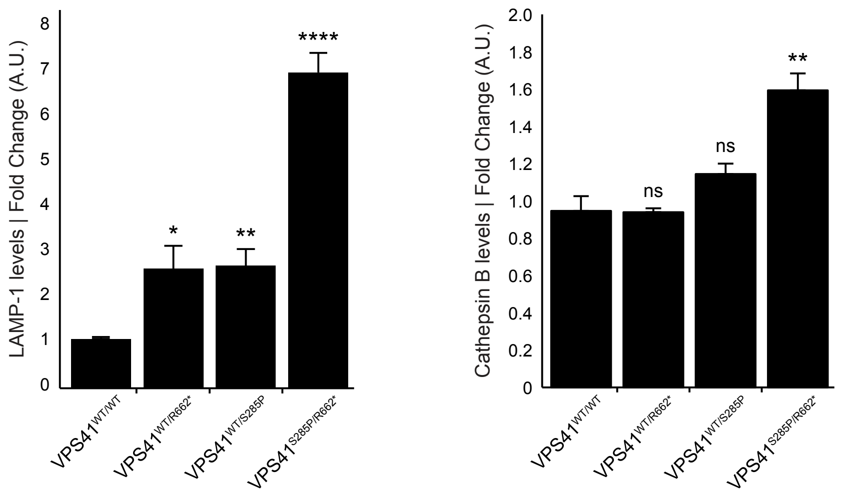


Figure S9

A

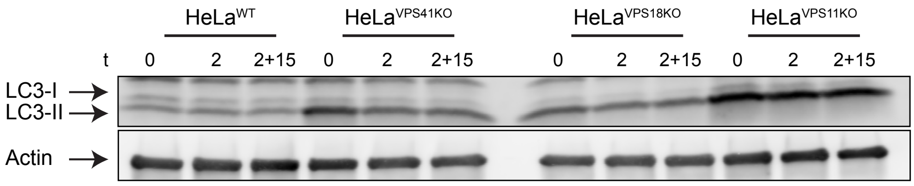


B

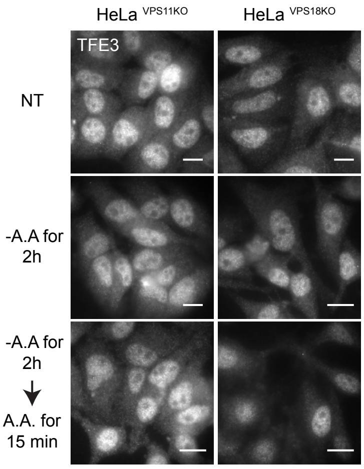


B´

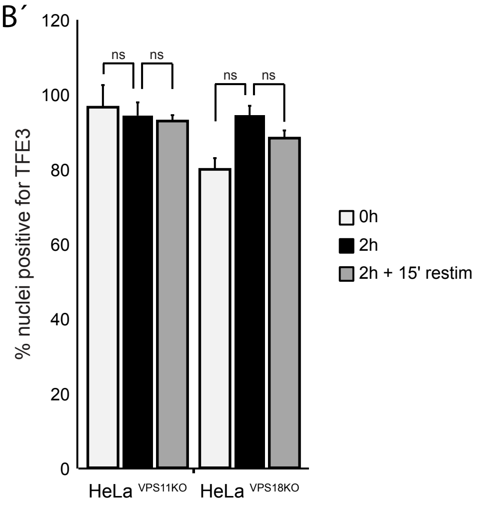


Figure S10

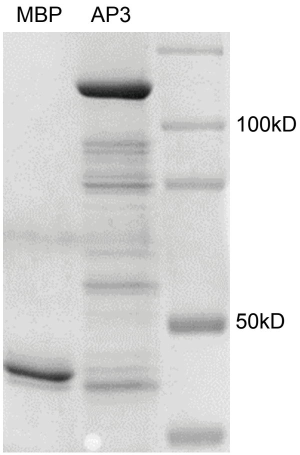

**Figure S8. Mutations in VPS41 results in increased levels of lysosomal proteins.** Western Blot analysis shows a significant increase in LAMP-1 and cathepsin B protein levels in *VPS41*$^{S285P/R662*}$ fibroblasts compared to *VPS41*$^{WT/WT}$, *VPS41*$^{WT/S285P}$ and *VPS41*$^{WT/R662*}$ fibroblasts (n=3). Error bars represent the SEM. *p<0.05, **p<0.01, ****p<10$^{-5}$; one-ANOVA with Bonferroni correction for multiple comparisons.

**Figure S9. Impaired HOPS complex results in impaired responsiveness to nutrient starvation. (A)** Western Blot of LC3-II to determine autophagosome formation in HeLa$^{VPS11KO}$, HeLa$^{VPS18KO}$ and HeLa$^{VPS41KO}$ cells. Increased LC3-II protein levels were observed for all 3 KO cell lines independent of nutrient availability. These results indicate that the autophagy phenotype is caused by impaired HOPS complex functionality (n=3). **(B)** HeLa$^{VPS11KO}$ and HeLa$^{VPS18KO}$ cells were either Nontreated (NT), starved for 2h (-AA) or starved and restimulated with nutrients (-AA (2h) + AA (15min)), fixed and labeled for TFE3. TFE3 is present in the nucleus of HeLa$^{VPS11KO}$ and HeLa$^{VPS18KO}$ cells regardless of nutrient availability, indicating nutrient insensitivity of these cells (quantified in **B'**). Error bars represent the SEM. (n=3); one-ANOVA with Bonferroni correction for multiple comparisons. Bars 10µm.

**Figure S10.** Representative coomassie stained gel of MBP and MBP-AP3 pulldown inputs.
